## Supporting Information for "TRAILS: tree reconstruction of ancestry using incomplete lineage sorting"

6 Model parameterization and optimization 24

7 Likelihood calculation and missing data 26

8 Implementation 27

9 Runtime 29

10 Parametric bootstrapping 30

11 Bibliography 30

### 1. INTRODUCTION

The main idea behind TRAILS is to extend the HMM in Mailund et al. [19] to accommodate one more species. The resulting model includes time discretization over a speciation tree with three species and an outgroup, while incorporating information about the topology of gene trees, similarly to CoalHMM [20, 21]. This supplementary material describes TRAILS in depth. Subsections 2.1 to 2.3 describe the continuous-time Markov chains with one, two and three lineages, respectively, necessary for the calculation of the transition probability matrix, while subsection 2.4 provides a short perspective on how TRAILS could be extended to accommodate more than three species. Section 3 provides an overview of the state space of the discrete-time Markov chain after dividing the coalescent space into discretized time intervals. Section 4 describes how to calculate the transition probability matrix, and section 5 explains how to compute the emission probabilities. Section 6 describes how the model is parameterized by population genetics parameters and how the optimization is performed, while section 7 shows how TRAILS handles missing data in the calculation of the likelihood. Finally, section 8 provides a short tutorial on how to run TRAILS in `python`.

### 2. CONTINUOUS-TIME MARKOV CHAINS

#### 2.1. Species A

Let A, B and C be present-day species, where, backwards in time, A and B are grouped first in the species tree, such that ((A,B),C); (fig. S1). In present day, all three species will have a sequence of nucleotides corresponding to a chromosome. We can approximate the coalescent with recombination of these samples by assuming that the process is Markovian, so the coalescent can be modelled using only two consecutive sites.

Focusing on species A, we will start with two linked sites, which are sampled at present. Going backwards in time, the two sites can stay linked, or, instead, they can recombine and sit in different chromosomes, each with their own genealogical history. After recombining, the two sites can coalesce back into the same chromosome. This coalescent with recombination process can be modelled using a one-sequence continuous-time Markov chain (CTMC), which has two possible states (fig. S2). The underlying transition rate matrix $\boldsymbol{Q}_{\mathbf{A}}$ summarizes the CTMC,

$$\boldsymbol{Q}_{\mathbf{A}} = \begin{pmatrix} -\gamma_{\mathrm{A}} & \gamma_{\mathrm{A}} \\ \rho_{\mathrm{A}} & -\rho_{\mathrm{A}} \end{pmatrix}, \tag{2}$$

where $\gamma_A$ and $\rho_A$ are the coalescent rate and the recombination rate in species A, respectively. In this CTMC, state 1 corresponds to the unlinked state, while state 2 represents the linked one (fig. S2). Equation (2) can also be visualized as a color-coded matrix, as showed in fig. S3. Grey indicates the diagonal entries, which are calculated as the negative of the sum of the off-diagonal entries in the corresponding row.

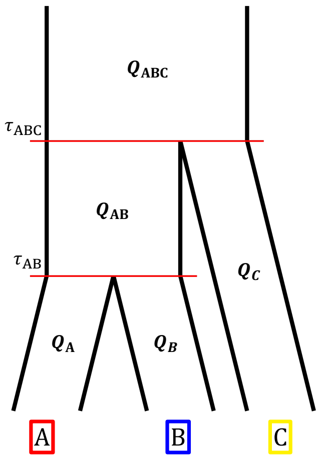

**Figure S1:** Species tree of the speciation process of three species (A, B and C). $\tau_{AB}$ and $\tau_{ABC}$ correspond to the speciation times, measured, backwards in time, as time since the present. The transition rate matrices for the CTMCs are also plotted in the part of the species tree that are used. These are three $2 \times 2$ matrices for the one-sequence CTMCs ($\boldsymbol{Q}_A, \boldsymbol{Q}_B, \boldsymbol{Q}_C$), a $15 \times 15$ matrix $\boldsymbol{Q}_{AB}$ for the two-sequence CTMC, and a $203 \times 203$ matrix $\boldsymbol{Q}_{ABC}$ for the three-sequence CTMC.

Following standard formulas for CTMCs, the probability of observing either of the two states of the one-sequence CTMC at time $\tau_{AB}$ can be calculated as the probability matrix $\exp(\tau_{AB}\boldsymbol{Q}_A)$. The first row of this $2 \times 2$ probability matrix represents the probability of being in each state at time $\tau_{AB}$ given that we start in state 1. Conversely, the second row represents the same, but given that the CTMC starts in state 2. Because the chain always starts in the linked state, the probabilities of being in each state at time $\tau_{AB}$ in the one-sequence CTMC are given by the vector $\boldsymbol{\pi}'_A = \boldsymbol{\pi}_A \exp(\tau_{AB}\boldsymbol{Q}_A)$, where $\boldsymbol{\pi}_A = (0, 1)$ is the vector of starting probabilities. Species B will follow a similar one-sequence CTMC, only with possibly different $\gamma_B$ and $\rho_B$, and transition rate matrix $\boldsymbol{Q}_B$. Thus, species B will also yield a vector of end probabilities $\boldsymbol{\pi}'_B$.

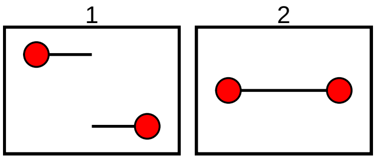

**Figure S2:** States of the CTMC of the coalescent-with-recombination process between two sites for a single sequence.

#### 2.2. Species AB

After reaching $\tau_{\mathrm{AB}}$, species A and B are mixed together. Forwards in time, this would correspond to a speciation event, where A and B become isolated. Backwards in time, the coalescent with recombination process can then be modelled using a two-sequence CTMC, similar to that described in Mailund et al. [19]. This CTMC has 15 different states (fig. S4), which correspond to all possible combinations of linked and recombined sequences for two left sites and two right sites. The corresponding transition rate matrix $\boldsymbol{Q}_{\mathbf{AB}}$ can be consulted in fig. S5, where $\gamma_{\mathrm{AB}}$ and $\rho_{\mathrm{AB}}$ are the coalescent rate and the recombination rate, respectively.

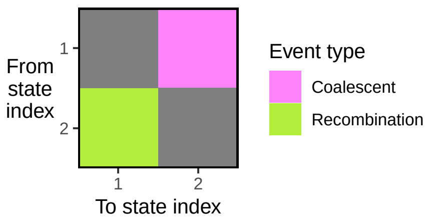

**Figure S3:** Transition rate matrix for the CTMC of the coalescent-with-recombination process between two sites for a single sequence. A coalescent event happens with rate $\gamma_A$ (in pink), while a recombination event happens with rate $\rho_A$ (in green).

Note that $\boldsymbol{Q}_{\mathbf{AB}}$ has a block-like structure, where some coalescent events prevent the return to previous states (see fig. S5). These events happen when two sites of different origin (one from A and one from B) reach their common ancestor. This type of coalescence is irreversible, in the sense that the two sequences coalesce and will now have a shared ancestral history. The other kind are coalescent events that place the left and right sites on the same chromosome, such as the one described for the one-sequence CTMC in subsection 2.1. This type of coalescence is reversible, meaning that recombination can separate the left and the right site into different chromosomes, thus unlinking their genetic history.

A way to more easily identify the blocks created by reversible and irreversible coalescent events is to define sets of states $\boldsymbol{\omega}$. If no irreversible coalescent has happened at either side (i.e. left and right sites), then that state belongs to the set $\boldsymbol{\omega}_{\mathbf{00}}$, where 0 represents no coalescent. If instead species A and B have coalesced at the left site, then that state belongs to the set $\boldsymbol{\omega}_{\mathbf{30}}$, and if species A and B have coalesced only at the right site, then that state belongs to $\boldsymbol{\omega}_{\mathbf{03}}$. The reason for the 3 subscript of $\boldsymbol{\omega}_{\mathbf{30}}$ and $\boldsymbol{\omega}_{\mathbf{03}}$ is that species A is assigned the number 1, while species B is assigned the number 2, so when these two coalesce, then $1 + 2 = 3$ (see fig. S6 for an explanatory diagram). Finally, if both the left and the right sites have coalesced, then that state belongs to $\boldsymbol{\omega}_{\mathbf{33}}$. In fact, the two states in $\boldsymbol{\omega}_{\mathbf{33}}$ are the absorbing states of this two-sequence CTMC (states 14 and 15 in fig. S4).

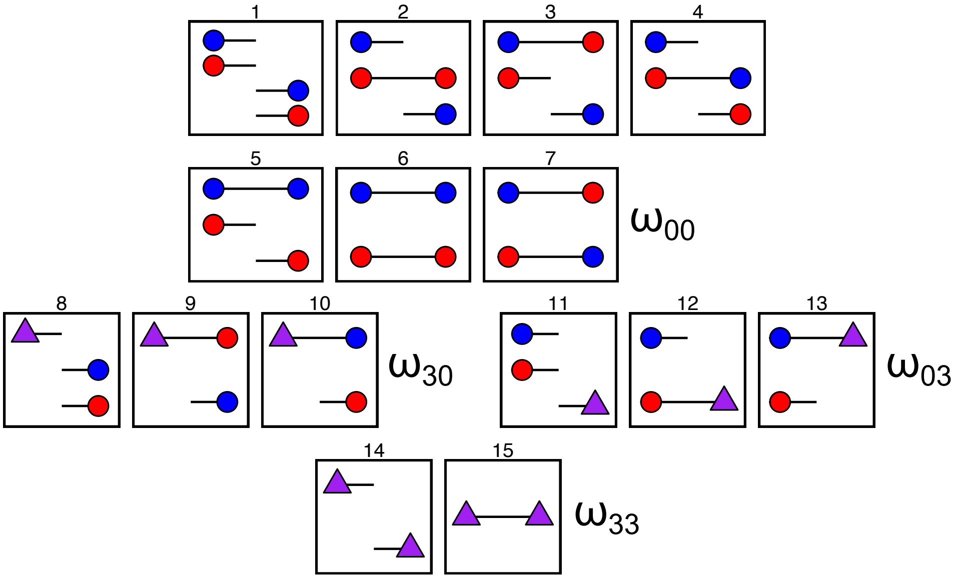

**Figure S4:** States of the CTMC of the coalescent-with-recombination process between two sites for two sequences. The states are grouped by $\boldsymbol{\omega}$ sets. Refer to fig. S6 for an explanation of the colors and shapes.

Similar to the one-sequence CTMC, the probability of observing each of the 15 states given that we start in either of them can be calculated using $\exp((\tau_{\text{ABC}} - \tau_{\text{AB}})\boldsymbol{Q}_{\mathbf{AB}})$. Now, unlike for the one-sequence CTMCs where the chain could only start from a single state, i.e. $\boldsymbol{\pi}_{\mathbf{A}} = (0, 1)$, we will have starting probabilities for each state $\boldsymbol{\pi}_{\mathbf{AB}}$ based on the mixing of the end probabilities of the two one-sequence CTMCs, $\boldsymbol{\pi}'_{\mathbf{A}}$ and $\boldsymbol{\pi}'_{\mathbf{B}}$. More specifically, $\boldsymbol{\pi}_{\mathbf{AB}}$ is calculated by multiplying the end probabilities each of the entries of the two one-sequence CTMCs. All in all, after running the two-sequence CTMC for a time of $\tau_{\text{ABC}} - \tau_{\text{AB}}$, the end probabilities can be calculated using $\boldsymbol{\pi}'_{\mathbf{AB}} = \boldsymbol{\pi}_{\mathbf{AB}} \exp((\tau_{\text{ABC}} - \tau_{\text{AB}})\boldsymbol{Q}_{\mathbf{AB}})$.

#### 2.3. Species ABC

Finally, the second speciation event is reached at time $\tau_{\text{ABC}}$. Backwards in time, this can be modelled as a three-sequence CTMC, containing 203 states (fig. S7). Again, these states correspond to all possible combinations among three linked and unlinked sequences. Similar to the two-sequence CTMC, the three-sequence case also includes irreversible coalescent events which generate a block-like structure in the transition rate matrix $\boldsymbol{Q}_{\mathbf{ABC}}$ (fig. S8). As with the two-sequence CTMC, these blocks define sets of states, here denoted $\boldsymbol{\Omega}$ with subscripts again specified as in fig. S6. The three-sequence CTMC also includes two absorbing states in the set $\boldsymbol{\Omega}_{77}$, where all three sequences have coalesced at both sides (states 202 and 203 in fig. S7 and fig. S8). All the $\boldsymbol{\Omega}$ sets and their correspondence to the states of the CTMC are shown in fig. S7.

The starting probability vector $\boldsymbol{\pi}_{\mathbf{ABC}}$ for the three-sequence CTMC can be calculated by mixing the

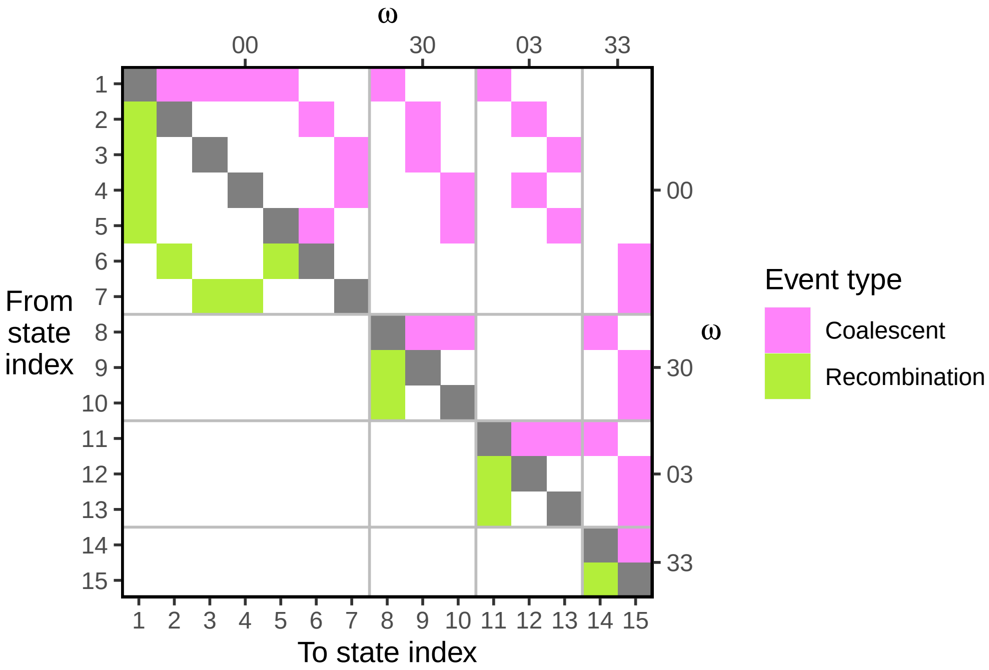

**Figure S5:** Transition rate matrix for the CTMC of the coalescent-with-recombination process between two sites for two sequences. A coalescent event happens with rate $\gamma_{AB}$ (in pink), while a recombination event happens with rate $\rho_{AB}$ (in green). The state indices in the left and bottom axes can be consulted in fig. S7. The right and top axes represent sets of states $\boldsymbol{\omega}$, which define the block-like structure of the transition rate matrix. Transitions with rates equal to zero are shown in white.

end probabilities of the two-sequence CTMC, or $\boldsymbol{\pi}'_{\mathbf{AB}}$, and the end probabilities of a one-sequence CTMC for the last species C, or $\boldsymbol{\pi}'_{\mathbf{C}}$. Similarly to how $\boldsymbol{\pi}'_{\mathbf{A}}$ and $\boldsymbol{\pi}'_{\mathbf{B}}$ are calculated, $\boldsymbol{\pi}'_{\mathbf{C}} = \boldsymbol{\pi}_{\mathbf{C}} \exp((\tau_{\mathrm{AB}} + \tau_{\mathrm{ABC}})\boldsymbol{Q}_{\mathrm{C}})$, where $\boldsymbol{\pi}_{\mathbf{C}} = (0, 1)$ is the vector of starting probabilities. The transition rate matrix $\boldsymbol{Q}_{\mathrm{C}}$ is parameterized

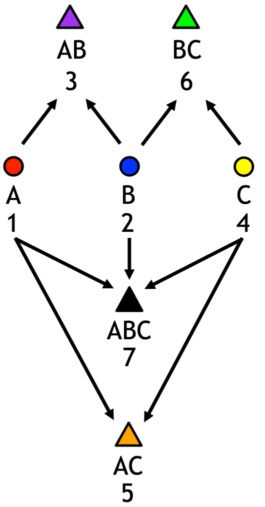

**Figure S6:** Diagram showing the different representations of sites used throughout this report. Colored circles represent uncoalesced sites, while triangles represent sites where at least one coalescent has happened. The numerical representation of each of the sites can be used to define the sets of states $\boldsymbol{\omega}$.

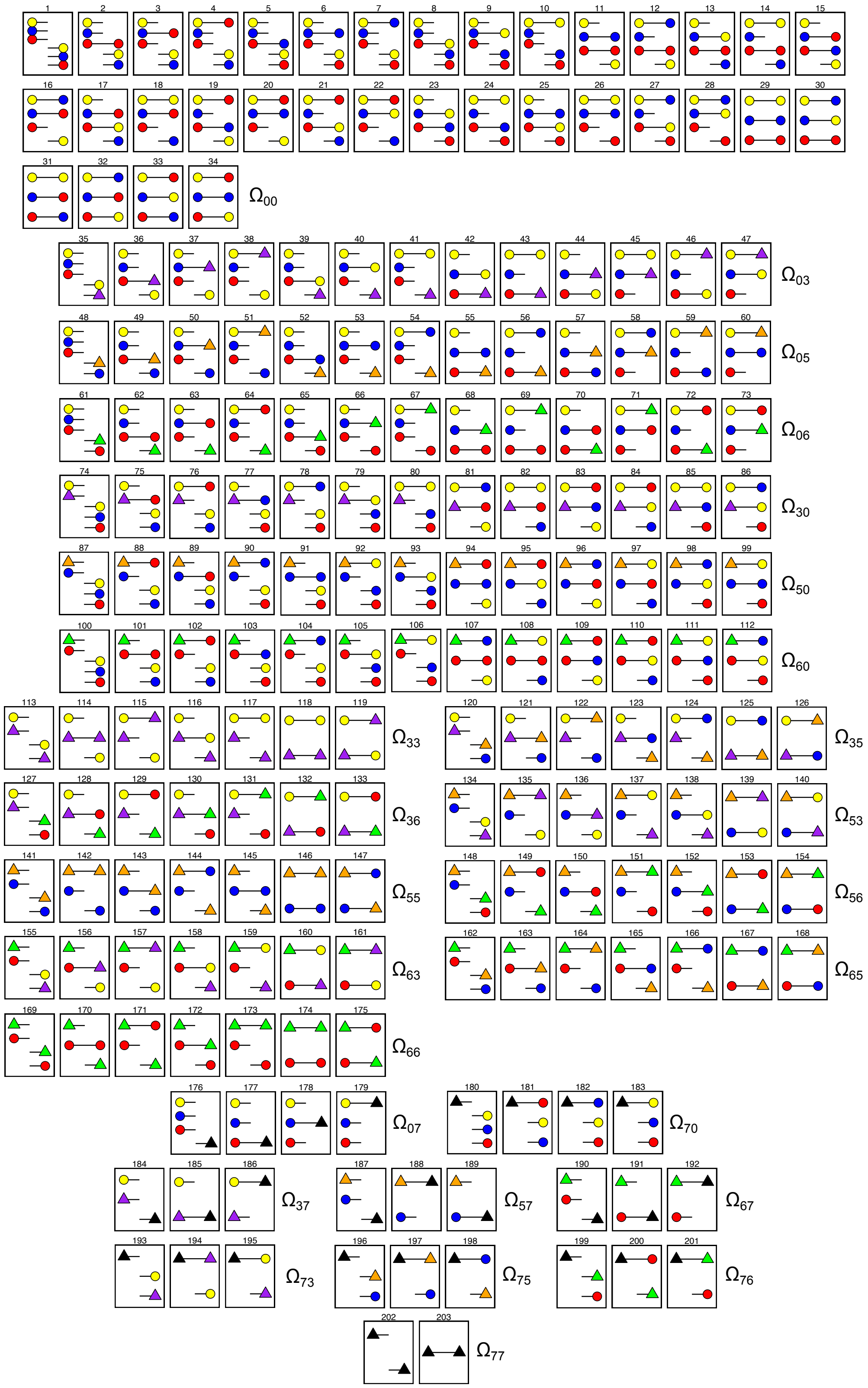

**Figure S7:** States of the CTMC of the coalescent-with-recombination process between two sites for three sequences. The states are grouped by $\mathbf{\Omega}$ sets. Refer to fig. S6 for an explanation of the colors and shapes.

by the coalescent rate $\gamma_C$ and the recombination rate $\rho_C$.

We can calculate the probability of observing any of the 203 states of the three-sequence CTMC at a given time point $t$ using $\boldsymbol{\pi}'_{\mathbf{ABC}}(t) = \boldsymbol{\pi}_{\mathbf{ABC}} \exp(t\boldsymbol{Q}_{\mathbf{ABC}})$. In any case, as $t$ tends to infinity, the probability vector will be concentrated on the two absorbing states of the CTMC, states 202 and 203 in which both sites have experienced two (irreversible) coalescent events and reached their most recent common ancestor.

It is of note to observe that the three-sequence CTMC contains states where only one coalescent has happened at either side. Focusing at a certain site, this first coalescent could have happened during the two-sequence CTMC, and, thus, the only possible tree is ((A,B),C). This tree follows the species tree

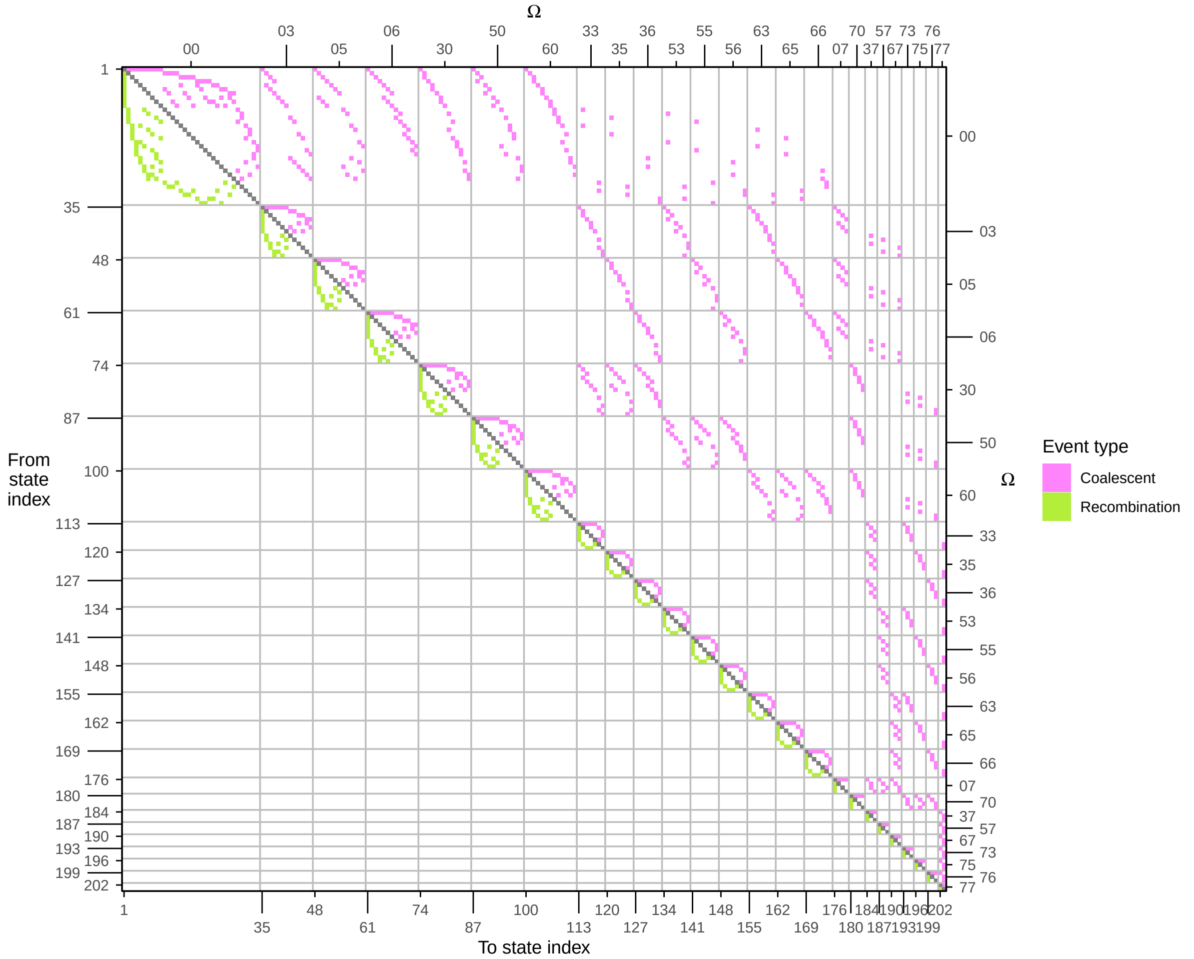

**Figure S8:** Transition rate matrix for the CTMC of the coalescent-with-recombination process between two sites for three sequences, or $\boldsymbol{Q}_{\mathbf{ABC}}$. A coalescent event happens with rate $\gamma_{ABC}$ (in pink), while a recombination event happens with rate $\rho_{ABC}$ (in green). The state indices in the left and bottom axes can be consulted in fig. S7. The right and top axes represent sets of states $\boldsymbol{\Omega}$, which define the block-like structure of the transition rate matrix. Again, transitions with rates equal to zero are shown in white.

(fig. S1), and it will be referred to as V0 (fig. S9). However, if no coalescence has happened at a site, then any pair of sequences can coalesce first, since sequences A, B and C are all mixed in the three-sequence CTMC. This means that trees such as ((A,C),B) or ((B,C),A) can also be generated in addition to ((A,B),C). These three trees of deep coalescent will be referred to as V1 if the tree follows the species tree topology (i.e. the ((A,B),C) tree), V2 for ((A,C),B), and V3 for ((B,C),A). V2 and V3 are incongruent with the species tree, meaning that their topologies are different from fig. S9. The proportion of sites that follow either of these two topologies is known as incomplete lineage sorting, or ILS for short.

ILS is a well-documented source of gene tree incongruence, and it is an obstacle for phylogenetic reconstruction. However, ILS also holds valuable information about the speciation process. In fact, the amount of ILS increases with the ancestral effective population size ($N_e$) between speciation events, and decreases the longer the time between speciation events is ($\tau_{ABC} - \tau_{AB}$). The exact formula for the expected proportion of ILS is $\Pr(\text{discordant topology}) = \frac{2}{3}\exp(-T/N_e)$, where $T = \tau_{ABC} - \tau_{AB}$ is measured in number of generations and $N_e$ is the haploid effective population size. Intuitively, if $N_e$ is large, then there is a higher chance that at least one site or lineage does not coalesce between the two speciation times. Moreover, if the speciation events happen in quick succession, then lineages will not have time to coalesce within the speciation times, and more sites will escape coalescing before $\tau_{ABC}$. Since ILS fragments are informative about the demography of the samples, modelling ILS enables the estimation of these population genetics parameters along the speciation process.

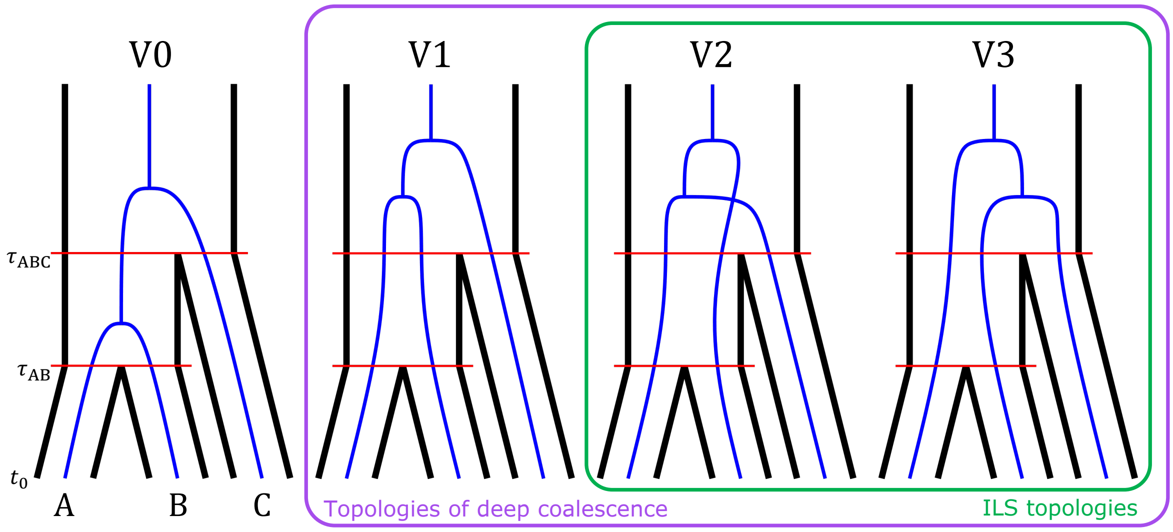

**Figure S9:** The four possible topologies that can arise during the speciation process of three species (A, B and C).

#### 2.4. Beyond three species

The procedure for building CTMCs is not in principle limited to three species. One could calculate the transition rate matrix for a CTMC with any number of lineages $n$. However, the number of states of the CTMC quickly explodes (see table S1), since it follows the even entries of the Bell number series. The number of states is given by the Bell numbers because each state can be thought of as a possible configuration of a set with $2n$ elements, corresponding to all sites and lineages involved in the CTMC. A group of such elements correspond to either the left and the right site being in the same chromosome, or to two sites at either the left or the right side having coalesced. The state space of the CTMC is then defined as all possible partitions of the elements. As a result of this, the state space quickly increases, and these large matrices can only be used directly for small values of $n$, since handling them often involve matrix inversions, multiplications and exponentiations, which are computationally intensive.

However, the information about the coalescent with recombination held in these matrices could potentially be mined with the help of some workarounds. For example, as shown in fig. S5 and fig. S8, the states of the rate matrices are organized in a block-like structure, and some of the states show symmetries. For specific applications, these large matrices can be reduced by merging states together. For example, for the two-sequence CTMC, we might be interested to drop the information about the origin of the lineages, and instead only keep track of whether there is a coalescent at either of the two sites, regardless of the labeling of the lineages. This is conceptually the same as removing the colors in fig. S4. In this situation, the state space is reduced from 15 states to just 8, by appropriately merging equivalent states and adjusting their transition rates. The same could be done with the states in the three-sequence CTMC (fig. S7), which would reduce the state space from 203 to just 31, or for a four-sequence CTMC, which would decrease the number of states from 4,140 to only 108. For the current application, however, the labeling of the lineages needs to be kept since it is important for modelling ILS, so the full state space is maintained.

Another property of these matrices is that they are extremely sparse, especially as $n$ increases. For example, 26.2% of the entries in the rate matrix for the two-sequence CTMC are non-zero (fig. S5). For the three-sequence CTMC (fig. S8), this proportion decreases to 3.2%, and, for a four-sequence CTMC, it is only 0.2%. Understanding the sparsity of these matrices could help reduce the computational burden by choosing the most optimal analytical procedures for costly operations such as matrix inversion or exponentiation.

In any case, if one is interested in obtaining the transition rate matrices for an arbitrary number of sequences, they can be computed using a similar approach as that described in this supplementary material. After enumerating the full state space using the Bell number series, we can encode each state in such a way that computing the valid transitions requires minimal bookkeeping. In this manuscript, we propose encoding the states using minimally superincreasing integers to represent all uncoalesced and coalesced sites with a unique numeric identifier (see fig. S6). One can then use an iterative approach to build the transition rate matrix by checking whether transitions through coalescence or recombination are allowed between pairs of states, and updating the numeric labels accordingly. We include an R script that employs this strategy to compute the transition rate matrix, which can be found in the accompanying GitHub repository under `https://github.com/rivasiker/trails_paper/tree/main/state_space_exploration`. We note, however, that the current implementation is only computationally feasible for a limited number of sequences ($n \leq 5$)

| **n** | **Number of states** | **Proportion of non-zero entries** |
|---|---|---|
| 1 | 2 | 100% |
| 2 | 15 | 26.2% |
| 3 | 203 | 3.2% |
| 4 | 4,140 | 0.2% |
| 5 | 115,975 | |
| 6 | 4,213,597 | |
| 7 | $1.91 \times 10^{8}$ | |
| 8 | $1.05 \times 10^{10}$ | |
| 9 | $6.82 \times 10^{11}$ | |
| 10 | $5.17 \times 10^{13}$ | |
| 100 | $6.25 \times 10^{275}$ | |
| 1000 | $1.24 \times 10^{4349}$ | |

**Table S1:** The number of states and proportion of non-zero entries in the rate matrix for a CTMC with $n$ lineages.

#### 3. The state space of the discrete-time Markov chain

As shown in the previous section, the speciation process can be described as a series of interconnected CTMCs. However, inferring population genetics parameters in continuous time is challenging. We can instead define a discrete-time Markov chain (DTMC), where the states correspond to three-leaved phylogenetic trees with two irreversible coalescent events. The coalescent times are allowed to happen at any point within the limits of distretized time intervals along the speciation tree. The transition probability matrix of this DTMC can then be used as the transition matrix of a hidden Markov model (HMM), where the observed states are the nucleotides of a three-way genome alignment (and an outgroup). The emission probabilities of the HMM will, then, be defined by the mutation rate and a nucleotide substitution model. This HMM can thus be used to estimate the underlying population genetic parameters along the speciation process. This sort of discretization is a common feature of applied coalescent HMMs. Note, this means that precise transition rates (which depend on the actual coalescence times) are replaced with bin-averaged transition rates. We expect better inferences if time is divided more finely, but adding more bins increases the computational burden.

The phylogenetic trees that constitute the states of the DTMC are conditioned by the path taken by the three sequences involved. For the sake of simplicity, let's assume that both the two-sequence CTMC and the three-sequence CTMC are divided into 3 discrete time intervals, or $n_{AB} = n_{ABC} = 3$. Then, let's imagine that, at a certain site, species A and species B coalesce within the 3rd interval of the two-sequence CTMC, and, later, that AB lineage coalesces with species C within the 2nd time interval of the three-sequence CTMC, generating a ((A,B),C) or V0 topology (fig. S10). This certain state of the DTMC will be referred to as $V0_3^2$. Generally, then, all possible combinations of V0 states can be referred to as $V0_x^y$, where the subscript $x$ represents the time interval in the two-sequence CTMC where the first coalescent has happened, and the superscript $y$ represents the time interval in the three-sequence CTMC where the second coalescent has happened. In total, there are $n_{AB} \times n_{ABC}$ different V0 states.

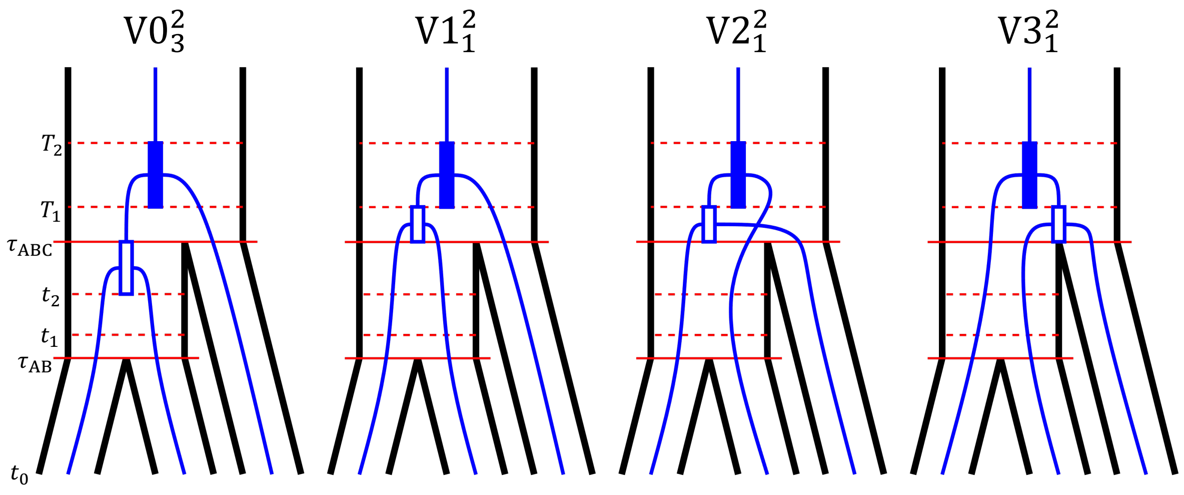

**Figure S10:** The hidden states of TRAILS. This figure shows four of the 27 possible hidden states for when $n_{AB} = n_{ABC} = 3$. In this figure, and in the rest of the figures of this report, the first coalescent is always plotted as an empty rectangle within an interval, meaning that the coalescent is allowed to happen at any point within the interval. The same goes for the second coalescent, which is in turn represented as a filled rectangle. Note that $t_0 = 0$.

Additionally, if the first coalescent does not happen between $\tau_{AB}$ and $\tau_{ABC}$, then both the first and the second coalescents must happen deep in the species tree during the three-sequence CTMC. If the tree still follows the ((A,B),C) or V1 tree, the first coalescent event happens in the 1st time interval of the three-sequence CTMC, and the second coalescent event happens in the 2nd time interval, then that state will be referred to as $V1_1^2$ (fig. S10). Again, more generally, all possible combinations of V1 states can be be represented by $V1_x^y$, where the subscript $x$ is the time interval in the three-sequence CTMC where the first coalescent has happened, and the superscript $y$ is the interval in the three-sequence

CTMC where the second coalescent has happened. Similarly, $V2_x^y$ and $V3_x^y$ also holds for all ((A,C),B) and ((B,C),A) trees, respectively (fig. S10). Note that sometimes $x = y$, so for the states of deep coalescent (V1, V2, and V3), both the first and the second coalescent can happen within the same time interval. Therefore, there are $n_{\text{ABC}} + \binom{n_{\text{ABC}}}{2}$ possible combinations for each deep coalescence state. All in all, there are $n_{\text{AB}} \times n_{\text{ABC}} + 3 \times \left[n_{\text{ABC}} + \binom{n_{\text{ABC}}}{2}\right]$ possible V0, V1, V2 and V3 states in total for the DTMC. As an example, all the 27 states for $n_{\text{AB}} = n_{\text{ABC}} = 3$ can be found in fig. S11.

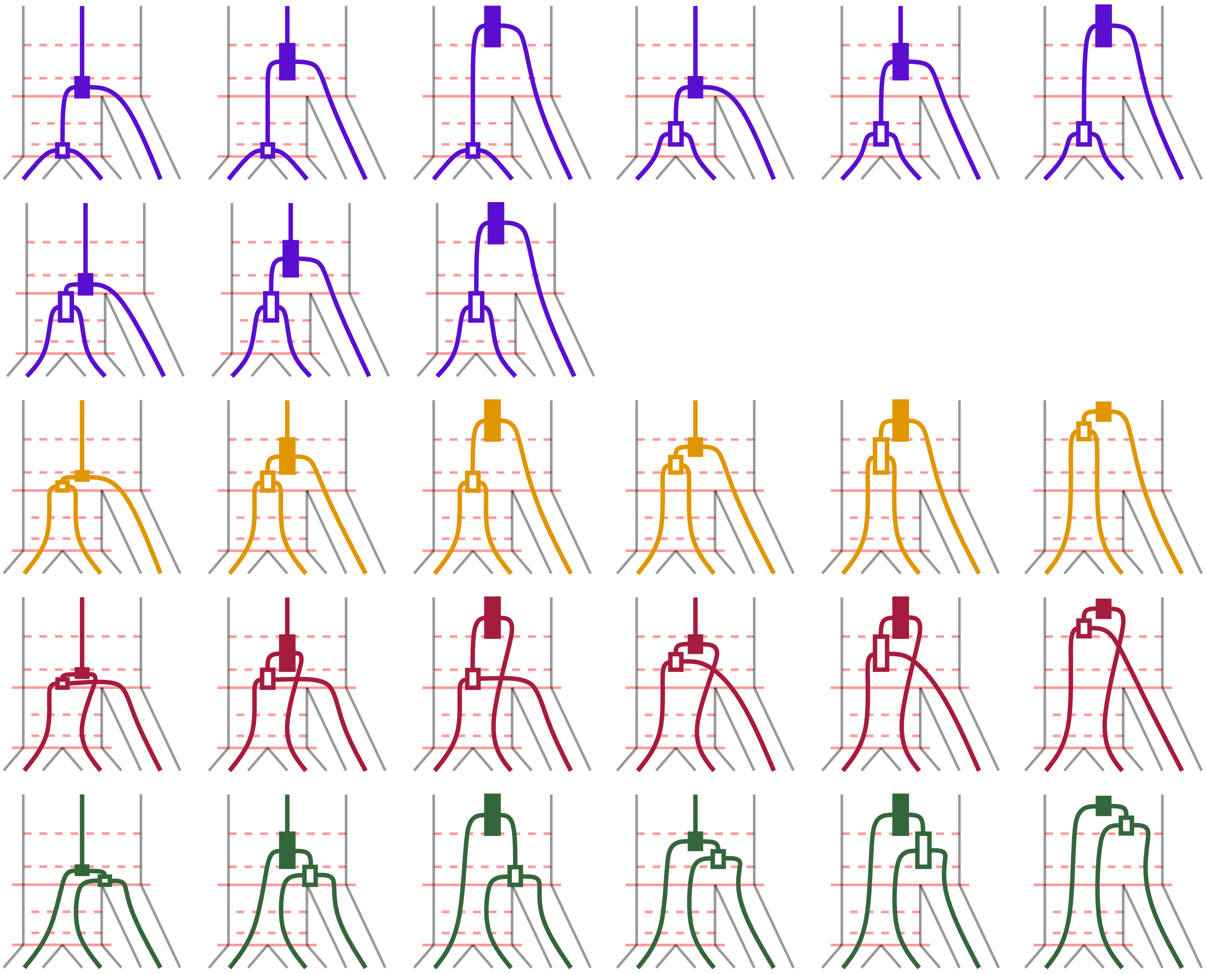

**Figure S11:** All 27 possible hidden states when $n_{\text{AB}} = n_{\text{ABC}} = 3$. The states are colored by topology.

### 4. THE TRANSITION PROBABILITY MATRIX

#### 4.1. The general case

To calculate the transition probability matrix of the DTMC, we need to consider the path taken by every pair of states, since each entry of the transition matrix is the probability of changing to the genealogy of the right-hand site R given that the left-hand site follows a certain genealogy L. Instead of calculating conditional probabilities, we can compute the joint probability matrix, where each entry is the joint probability of the left-hand site following a L genealogy, and the right-hand site following a R genealogy, or $\Pr(\mathrm{L}, \mathrm{R})$. In fact, we can easily convert the joint probability to conditional probability and, thus, to the probabilities in the transition matrix, knowing that $\Pr(\mathrm{L}, \mathrm{R}) = \Pr(\mathrm{L}) \times \Pr(\mathrm{R} \mid \mathrm{L})$.

In general, the joint probability can be calculated by obtaining the probability of observing either of the two absorbing states of the three-sequence CTMC ($\boldsymbol{\Omega_{77}}$) conditioned on the path taken, i.e. conditioned on L and R. This can generally be calculated using matrix exponentiation and choosing the states that are allowed at the beginning and end of each discretized time interval.

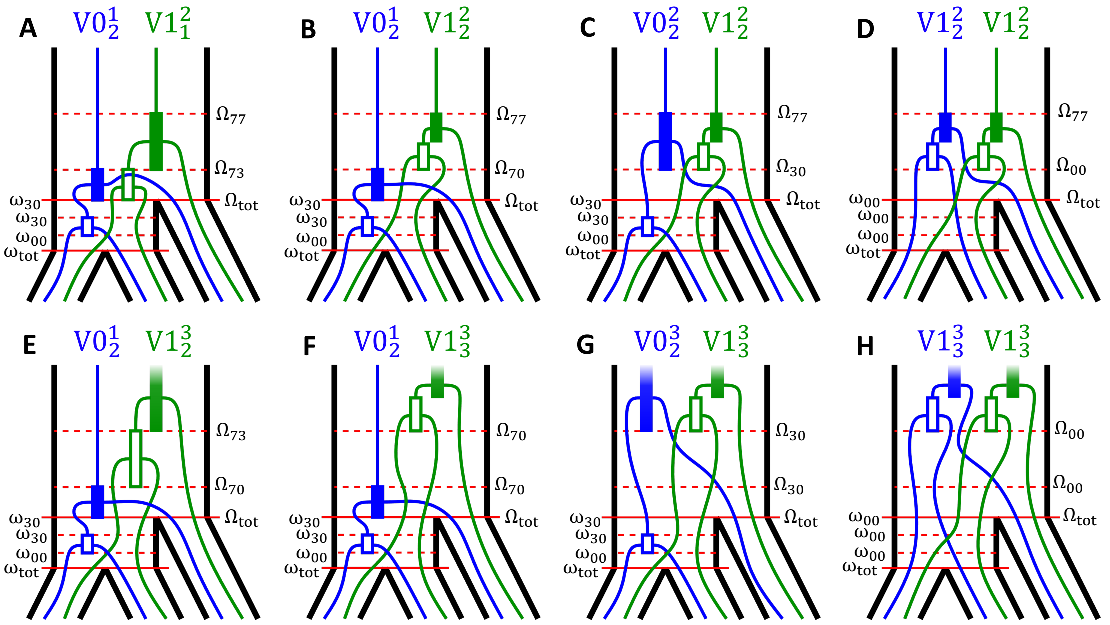

**Figure S12:** Paths and transitions. This figure shows 8 of the 729 possible combinations of L and R when $n_{\mathrm{AB}} = n_{\mathrm{ABC}} = 3$. The lowercase omegas ($\omega$) at the left-hand side of each tree show the set of possible states that are allowed at the beginning and end of each time interval in the two-sequence CTMC, while the uppercase omegas ($\Omega$) at the right-hand side show the same for the three-sequence CTMC. The names for the interval limits marked by red horizontal lines can be consulted in fig. S10.

As an example, let's assume that $n_{\mathrm{AB}} = 3$ and $n_{\mathrm{ABC}} = 3$, and that the left site follows a $\mathrm{L} = \mathrm{V0}^1_2$ tree, while the right site follows a $\mathrm{R} = \mathrm{V1}^2_1$ tree (fig. S12A). If we focus first on the two-sequence CTMC between the two speciation events, we can calculate the probability matrix at the end of the 1st interval as $\exp((t_1 - \tau_{\mathrm{AB}})\boldsymbol{Q_{\mathrm{AB}}})$. However, given the fact that neither the left site nor the right site have coalesced within this first interval, only a subset of the states of the CTMC are allowed, namely those where no coalescent has happened at either side. Thus, we can get a sub-matrix by choosing the right row and column indices of $\exp((t_1 - \tau_{\mathrm{AB}})\boldsymbol{Q_{\mathrm{AB}}})$, such that $\exp((t_1 - \tau_{\mathrm{AB}})\boldsymbol{Q_{\mathrm{AB}}})\,[\boldsymbol{\omega_{\mathrm{tot}}}; \boldsymbol{\omega_{00}}]$, where $\boldsymbol{\omega_{\mathrm{tot}}} = (1, 2, ..., 15)$ indicates that the CTMC can start from any of the 15 states, and $\boldsymbol{\omega_{00}}$ is a vector of indices of the 7 states where no coalescence has happened yet, which correspond to the states where the

CTMC can finish at the end of the 1st interval. Additionally, $t_1$ is the time at which the 1st interval ends (refer to fig. S10 for the interval times). The first and second vectors inside square brackets following a matrix represent, respectively, the row indices and the column indices to be kept for a desired sub-matrix.

Afterwards, the left site coalesces in the 2nd interval, while the right site remains uncoalesced. In this case, the probability sub-matrix will be $\exp((t_2 - t_1)\boldsymbol{Q}_{\mathbf{AB}})\,[\boldsymbol{\omega}_{\mathbf{00}}; \boldsymbol{\omega}_{\mathbf{30}}]$, where $\boldsymbol{\omega}_{\mathbf{30}}$ is a vector containing the indices of the CTMC states where the coalescent has happened on the left-hand site, and $t_2$ is the time at which the 2nd interval ends. The 30 subscript of $\boldsymbol{\omega}_{\mathbf{30}}$ means that species A (tagged with the number 1) and species B (tagged with the number 2) coalesce, so then $1 + 2 = 3$ at the left-hand site (see the diagram in fig. S6). A subscript of 0 means that no coalescent has happened yet, in this case referring to the right-hand site.

Similarly, since no coalescent events happen within the 3rd interval, the probability sub-matrix for this interval can be calculated as $\exp((\tau_{\text{ABC}} - t_2)\boldsymbol{Q}_{\mathbf{AB}})\,[\boldsymbol{\omega}_{\mathbf{30}}; \boldsymbol{\omega}_{\mathbf{30}}]$. All in all, in order to get the final probability sub-matrix $\boldsymbol{P}_{\mathbf{AB}}(\text{L}, \text{R})$ of observing a certain $\boldsymbol{\omega}_{\mathbf{30}}$ state at time $\tau_{\text{ABC}}$ given that we start in each of the states $\boldsymbol{\omega}_{\mathbf{tot}}$, and assuming that the left and right sites follow a $\text{L} = \text{V}0_2^1$ and $\text{R} = \text{V}1_1^2$ tree, respectively, we need to multiply all the probability sub-matrices. Thus,

$$
\begin{aligned}
&\boldsymbol{P}_{\mathbf{AB}}(\text{L} = \text{V}0_2^1, \text{R} = \text{V}1_1^2) = \\
&= e^{(t_1 - \tau_{\text{AB}})\boldsymbol{Q}_{\mathbf{AB}}}\,[\boldsymbol{\omega}_{\mathbf{tot}}; \boldsymbol{\omega}_{\mathbf{00}}]\; e^{(t_2 - t_1)\boldsymbol{Q}_{\mathbf{AB}}}\,[\boldsymbol{\omega}_{\mathbf{00}}; \boldsymbol{\omega}_{\mathbf{30}}]\; e^{(\tau_{\text{ABC}} - t_2)\boldsymbol{Q}_{\mathbf{AB}}}\,[\boldsymbol{\omega}_{\mathbf{30}}; \boldsymbol{\omega}_{\mathbf{30}}]\,,
\end{aligned}
$$

which is a $|\boldsymbol{\omega}_{\mathbf{tot}}| \times |\boldsymbol{\omega}_{\mathbf{30}}|$ probability sub-matrix. In a more general way,

$$
\boldsymbol{Z}(\boldsymbol{Q}, \boldsymbol{t}, \boldsymbol{\omega}) = \prod_{i=1}^{n-1} e^{(t_{i+1} - t_i)\boldsymbol{Q}}\,[\boldsymbol{\omega}_{\boldsymbol{i}}; \boldsymbol{\omega}_{\boldsymbol{i+1}}]\,, \tag{3}
$$

where $\boldsymbol{Q}$ is the transition rate matrix, $\boldsymbol{t}$ is the vector of time intervals of size $n$, and $\boldsymbol{\omega}$ is the vector of $n$ vectors of state indices for each of the interval cutpoints given a certain path for the left and the right sites. For our specific example, $\boldsymbol{P}_{\mathbf{AB}}(\text{L} = \text{V}0_2^1, \text{R} = \text{V}1_1^2) = \boldsymbol{Z}(\boldsymbol{Q}, \boldsymbol{t}, \boldsymbol{\omega})$, where $\boldsymbol{Q} = \boldsymbol{Q}_{\mathbf{AB}}$, $\boldsymbol{t} = (\tau_{\text{AB}}, t_1, t_2, \tau_{\text{ABC}})$ and $\boldsymbol{\omega} = (\boldsymbol{\omega}_{\mathbf{tot}}, \boldsymbol{\omega}_{\mathbf{00}}, \boldsymbol{\omega}_{\mathbf{30}}, \boldsymbol{\omega}_{\mathbf{30}})$ (refer to fig. S4 to consult exactly which states are included in each of the elements of $\boldsymbol{\omega}$). By careful choice of $\boldsymbol{\omega}$, we can calculate $\boldsymbol{P}_{\mathbf{AB}}(\text{L}, \text{R})$ for any combination of L and R using eq. (3).

In order to obtain the end probability vector for each combination of L and R, namely $\boldsymbol{\pi}'_{\mathbf{AB}}(\text{L}, \text{R})$, we need to multiply the starting probability vector of the two-sequence CTMC, $\boldsymbol{\pi}_{\mathbf{AB}}$, with the probability matrix, such that $\boldsymbol{\pi}'_{\mathbf{AB}}(\text{L}, \text{R}) = \boldsymbol{\pi}_{\mathbf{AB}}\boldsymbol{P}_{\mathbf{AB}}(\text{L}, \text{R})$. As described above in subsection 2.1, we can calculate the starting probability vector of the two-sequence CTMC ($\boldsymbol{\pi}_{\mathbf{AB}}$) by running one one-sequence CTMCs per species A and B, and combining their end probabilities.

Similar to the two-sequence CTMC, we can also use eq. (3) for the three-sequence case, such that the probability sub-matrix for a certain combination of L and R follows $\boldsymbol{P}_{\mathbf{ABC}}(\mathrm{L},\mathrm{R}) = \boldsymbol{Z}(\boldsymbol{Q},\boldsymbol{t},\boldsymbol{\omega})$. Keeping the example above, namely when $\mathrm{L} = \mathrm{V0}_2^1$ and $\mathrm{R} = \mathrm{V1}_1^2$ (see fig. S12A), $\boldsymbol{Q} = \boldsymbol{Q}_{\mathbf{ABC}}$, which is the $203 \times 203$ transition rate matrix of the two-sequence CTMC, $\boldsymbol{t} = (\tau_{\mathrm{ABC}}, T_1, T_2)$, where $T_1$ and $T_2$ are the end times of the 1st and 2nd discretized time intervals (see fig. S10 for the times), and $\boldsymbol{\omega} = (\boldsymbol{\Omega}_{\mathbf{tot}}, \boldsymbol{\Omega}_{\mathbf{33}}, \boldsymbol{\Omega}_{\mathbf{77}})$. For the latter, $\boldsymbol{\Omega}_{\mathbf{tot}} = (1,2,...,203)$, which represents the whole state space of the three-sequence CTMC, $\boldsymbol{\Omega}_{\mathbf{33}}$ is the vector of indices of possible end states after the 1st time interval, where only states where sequences A and B have coalesced at both the left and the right sides are allowed, and $\boldsymbol{\Omega}_{\mathbf{77}}$ is the vector of size 2 corresponding to the two absorbing states, which are the states where all three sequences have coalesced at both sites (refer to fig. S7 to consult which states are included in each $\boldsymbol{\Omega}$ set).

The joint probability of $\mathrm{L} = \mathrm{V0}_2^1$ and $\mathrm{R} = \mathrm{V1}_1^2$ can then be calculated as $\Pr(\mathrm{L} = \mathrm{V0}_2^1, \mathrm{R} = \mathrm{V1}_1^2) = \boldsymbol{\pi}_{\mathbf{ABC}}\boldsymbol{P}_{\mathbf{ABC}}(\mathrm{L},\mathrm{R})\boldsymbol{e}$, where $\boldsymbol{\pi}_{\mathbf{ABC}}$ is a vector of size 203 with the starting probabilities of the three-sequence CTMC obtained by mixing the end probabilities of the two-sequence CTMC ($\boldsymbol{\pi}'_{\mathbf{AB}}$) and the end probabilities of the one-sequence CTMC for species C ($\boldsymbol{\pi}'_{\mathbf{C}}$). $\boldsymbol{e}$ is simply a column vector of ones of size 2, i.e. $(1,1)^T$, signifying that the joint probability is the sum of being in either of the absorbing states at the end of the appropriate interval.

#### 4.2. Multiple coalescents within the same time interval

In many cases, $\Pr(\mathrm{L},\mathrm{R}) = \boldsymbol{\pi}_{\mathbf{ABC}}\boldsymbol{P}_{\mathbf{ABC}}(\mathrm{L},\mathrm{R})\boldsymbol{e}$, as described above. However, some combinations of L and R require further calculations. For example, when $n_{\mathrm{AB}} = n_{\mathrm{ABC}} = 3$, if $\mathrm{L} = \mathrm{V0}_2^1$ and $\mathrm{R} = \mathrm{V1}_2^2$, the first and the second coalescents of the right site happen within the same discretized time interval (fig. S12B). When this is the case,

$$\Pr(\mathrm{L} = \mathrm{V0}_2^1, \mathrm{R} = \mathrm{V1}_2^2) =$$

$$\boldsymbol{\pi}_{\mathbf{ABC}}(\mathrm{L},\mathrm{R})\; e^{(\boldsymbol{T_1}-\tau_{\mathrm{ABC}})\boldsymbol{Q}_{\mathrm{ABC}}}\,[\boldsymbol{\Omega}_{\mathbf{tot}}; \boldsymbol{\Omega}_{\mathbf{70}}] \int_{T_1}^{T_2} e^{(\boldsymbol{s}-\boldsymbol{T_1})\boldsymbol{Q}_{\mathrm{ABC}}} \boldsymbol{A}_{\mathbf{70,73}} e^{(\boldsymbol{T_2}-\boldsymbol{s})\boldsymbol{Q}_{\mathrm{ABC}}}\, ds\, [\boldsymbol{\Omega}_{\mathbf{70}}; \boldsymbol{\Omega}_{\mathbf{77}}]\; \boldsymbol{e},$$

where $\boldsymbol{A}_{\mathbf{70,73}}$ is a rate matrix of instantaneous transitions. This means that all entries in $\boldsymbol{A}_{\mathbf{70,73}}$ where the starting state is within $\boldsymbol{\Omega}_{\mathbf{70}}$, and the end state is within $\boldsymbol{\Omega}_{\mathbf{73}}$ equal to the value in the corresponding entry in $\boldsymbol{Q}_{\mathbf{ABC}}$, while all other values equal to 0.

The integral of the equation above can be calculated using Van Loan's approach [81]. Let $\boldsymbol{G}(t) =$

$\int_0^t e^{(t-s)\boldsymbol{Q}} \boldsymbol{A} e^{s\boldsymbol{Q}}\, ds$. We can construct a matrix $\boldsymbol{C}$, such that

$$\boldsymbol{C} = \begin{pmatrix} \boldsymbol{Q} & \boldsymbol{A} \\ \boldsymbol{0} & \boldsymbol{Q} \end{pmatrix}, \tag{4}$$

where $\boldsymbol{0}$ is a matrix of zeros of the same size as $\boldsymbol{Q}$ (or $\boldsymbol{A}$). Following Van Loan [81],

$$e^{t\boldsymbol{C}} = \begin{pmatrix} \boldsymbol{F}(t) & \boldsymbol{G}(t) \\ \boldsymbol{0} & \boldsymbol{F}(t) \end{pmatrix}, \tag{5}$$

where $\boldsymbol{F}(t) = e^{t\boldsymbol{Q}}$. Therefore, we can compute the matrix integral of interest, $\boldsymbol{G}(t)$, by taking the sub-matrix of size $|\boldsymbol{Q}|$ in the upper right corner of $e^{t\boldsymbol{C}}$.

Van Loan's method is also useful for calculating the joint probability of the left and the right topology when more than two coalescents happen within the same time interval. This can happen, for example, when $n_{\text{AB}} = n_{\text{ABC}} = 3$, $\text{L} = \text{V}0_2^2$ and $\text{R} = \text{V}1_2^2$, such that the 2nd coalescent at the left site and both the 1st and the 2nd coalescents at the right site happen within the 2nd interval of the three-sequence CTMC (fig. S12C). In this case, the integral will be of the form

$$\boldsymbol{G}(t) = \int_0^t \int_0^s e^{(t-s)\boldsymbol{Q}} \boldsymbol{A_1} e^{(s-r)\boldsymbol{Q}} \boldsymbol{A_2} e^{r\boldsymbol{Q}}\, dr\, ds,$$

which can also be calculated following Van Loan [81]. In fact, this same method can also be used to calculate triple integrals of the form

$$\boldsymbol{G}(t) = \int_0^t \int_0^s \int_0^r e^{(t-s)\boldsymbol{Q}} \boldsymbol{A_1} e^{(s-r)\boldsymbol{Q}} \boldsymbol{A_2} e^{(r-w)\boldsymbol{Q}} \boldsymbol{A_3} e^{w\boldsymbol{Q}}\, dw\, dr\, ds,$$

that arise when all four coalescents happen within the same interval, for example, when $\text{L} = \text{R} = \text{V}1_2^2$ (fig. S12D).

For some combinations of L and R, there is an additional level of complexity, since several paths need to be considered. For example, when $n_{\text{AB}} = n_{\text{ABC}} = 3$, $\text{L} = \text{V}0_2^2$ and $\text{R} = \text{V}1_2^2$ (fig. S12C), the integral part needs to consider 5 different paths:

- $30 \rightarrow 33 \rightarrow 73 \rightarrow 77$
- $30 \rightarrow 33 \rightarrow 37 \rightarrow 77$
- $30 \rightarrow 70 \rightarrow 73 \rightarrow 77$
- $30 \rightarrow 73 \rightarrow 77$

- $30 \rightarrow 33 \rightarrow 77$

These can be computed separately, and then summed over to calculate the joint probability of L and R. The number of paths to be considered can quickly increase, especially when all coalescents happen in the same time interval. The current implementation of the algorithm uses all available cores to parallelize these calculations.

#### 4.3. The deepest time interval

All the equations described above can only be used when the intervals when coalescents happen have finite time limits. This poses a problem when coalescents happen within the last time interval, since its upper limit is infinity. A workaround can be devised by knowing that the CTMC is absorbing, so once the last interval is reached, absorption within the interval is guaranteed to occur.

If a single coalescent event happens within the last interval, as is the case for $\mathrm{L} = \mathrm{V0}_2^1$ and $\mathrm{R} = \mathrm{V1}_2^3$ when $n_{\mathrm{AB}} = n_{\mathrm{ABC}} = 3$ (fig. S12E), then $\Pr(\mathrm{L}, \mathrm{R}) = \boldsymbol{\pi}_{\mathbf{ABC}} \boldsymbol{Z}(\boldsymbol{Q}, \boldsymbol{t}, \boldsymbol{\omega}) \boldsymbol{e}$, where $\boldsymbol{Q} = \boldsymbol{Q}_{\mathbf{ABC}}$, $\boldsymbol{t} = (\tau_{\mathrm{ABC}}, T_1, T_2)$, $\boldsymbol{\omega} = (\boldsymbol{\Omega}_{\mathbf{tot}}, \boldsymbol{\Omega}_{\mathbf{70}}, \boldsymbol{\Omega}_{\mathbf{73}})$, and $\boldsymbol{e}$ is a column vector of ones of size $|\boldsymbol{\Omega}_{\mathbf{73}}|$. In other words, we avoid the last interval by knowing that all states that have coalesced in previous intervals to any of the $\boldsymbol{\Omega}_{\mathbf{73}}$ states will eventually coalesce to one of the two absorbing states $\boldsymbol{\Omega}_{\mathbf{77}}$ within the last interval given enough time, since that is the only path they can take. We can also arrive to this conclusion by looking at the block structure of $\boldsymbol{Q}_{\mathbf{ABC}}$ (fig. S8).

If, instead, two coalescent events happen within the last interval, then we need to calculate integrals of matrix exponentials with an infinite upper limit. For example, when $n_{\mathrm{AB}} = n_{\mathrm{ABC}} = 3$, $\mathrm{L} = \mathrm{V0}_2^1$ and $\mathrm{R} = \mathrm{V1}_3^3$ (fig. S12F),

$$\Pr(\mathrm{L}, \mathrm{R}) = \pi_{\mathrm{ABC}} \boldsymbol{Z}(\boldsymbol{Q}, \boldsymbol{t}, \boldsymbol{\omega}) \int_0^\infty e^{(t-s)Q_{\mathrm{ABC}}} \boldsymbol{A}_{\mathbf{70,73}} \, ds \, [\boldsymbol{\Omega}_{\mathbf{70}}; \boldsymbol{\Omega}_{\mathbf{73}}] \; \boldsymbol{e},$$

where $\boldsymbol{Q} = \boldsymbol{Q}_{\mathbf{ABC}}$, $\boldsymbol{t} = (\tau_{\mathrm{ABC}}, T_1, T_2)$, $\boldsymbol{\omega} = (\boldsymbol{\Omega}_{\mathbf{tot}}, \boldsymbol{\Omega}_{\mathbf{70}}, \boldsymbol{\Omega}_{\mathbf{70}})$, and $\boldsymbol{e}$ is a column vector of ones of size $|\boldsymbol{\Omega}_{\mathbf{73}}|$. Looking at the integral, we can see it is of the form

$$\int_0^\infty e^{s\boldsymbol{Q}} \boldsymbol{A} \, ds = \int_0^\infty e^{s\boldsymbol{Q}} \, ds \boldsymbol{A} = -(\boldsymbol{Q})^{-1} \boldsymbol{A}.$$

This equation cannot be computed if $\boldsymbol{Q} = \boldsymbol{Q}_{\mathbf{ABC}}$, because $\boldsymbol{Q}_{\mathbf{ABC}}$ is a singular matrix. However, we can compute it if $\boldsymbol{Q} = \boldsymbol{Q}'_{\mathbf{ABC}}$, where $\boldsymbol{Q}'_{\mathbf{ABC}}$ is a sub-matrix of $\boldsymbol{Q}_{\mathbf{ABC}}$ without the two absorbing states. Using this sub-intensity matrix makes sense, since, in fact, we are only interested in the coalescents that have happened before the last coalescent, and all those states will eventually coalesce to the absorbing states given enough time.

If instead of two there are three coalescents happening on the last interval, as is the case for L $= \mathrm{V0}_2^3$ and R $= \mathrm{V1}_3^3$ when $n_{\mathrm{AB}} = n_{\mathrm{ABC}} = 3$ (fig. S12G), then we will need to compute a double integral of the form

$$\int_0^\infty \int_0^s e^{(s-r)Q} \boldsymbol{A_1} e^{rQ} \boldsymbol{A_2} \, dr \, ds = \int_0^\infty \int_0^s e^{(s-r)Q} \boldsymbol{A_1} e^{rQ} \, dr \, ds \boldsymbol{A_2} = \int_0^\infty \boldsymbol{G}(s) \, ds \boldsymbol{A_2}.$$

In order to compute $\boldsymbol{G}(s)$, we can use [81] by calculating $\boldsymbol{C}$ following eq. (4), where $\boldsymbol{A} = \boldsymbol{A_1}$ and $\boldsymbol{Q} = \boldsymbol{Q'_{\mathrm{ABC}}}$. Then, following eq. (5), we obtain a matrix exponential $\exp(t\boldsymbol{C})$ that depends on time. Integrating such exponential, we get

$$\int_0^\infty e^{s\boldsymbol{C}} \, ds = -(\boldsymbol{C})^{-1} = \begin{pmatrix} \boldsymbol{F} & \int_0^\infty \boldsymbol{G}(s) \, ds \\ \boldsymbol{0} & \boldsymbol{F} \end{pmatrix},$$

where its upper-right sub-matrix of size $|\boldsymbol{Q'_{\mathrm{ABC}}}|$ equals to the integral of interest, and $\boldsymbol{F} = -(\boldsymbol{A_1})^{-1}$. Following a similar procedure, we can also compute the triple integral needed when all coalescents happen within the last interval, for example, when $n_{\mathrm{AB}} = n_{\mathrm{ABC}} = 3$ and L = R $= \mathrm{V1}_3^3$ (fig. S12H). These integrals are of the form

$$\int_0^\infty \int_0^s \int_0^r e^{(s-r)Q} \boldsymbol{A_1} e^{(r-w)Q} \boldsymbol{A_2} e^{wQ} \boldsymbol{A_3} \, dw \, dr \, ds =$$
$$= \int_0^\infty \int_0^s \int_0^r e^{(s-r)Q} \boldsymbol{A_1} e^{(r-w)Q} \boldsymbol{A_2} e^{wQ} \, dw \, dr \, ds \boldsymbol{A_3} = \int_0^\infty \boldsymbol{G}(s) \, ds \boldsymbol{A_3}.$$

Note that, as explained in the previous section, some combinations of L and R will have more than one possible path, so in order to compute their joint probability, we need to calculate the probability of each path separately, and then sum these probabilities.

### 5. THE EMISSION PROBABILITIES

#### 5.1. The observed states

The states of the DTMC describe the marginal genealogical histories of the sequences. However, these states cannot be observed directly. Instead, each of the states will produce site patterns with different probabilities based on the genealogy they follow. Thus, the states of the DTMC can be used as the latent states of a hidden Markov model (HMM), and the nucleotide patterns as the observed states. The observed states will thus correspond to columns along a 4-way genome alignment, with three species and an outgroup. In the absence of missing data, the number of possible observed states can be calculated as the number of combinations of the four possible nucleotides (A, C, G, T) for four sequences, which amounts to $4^4 = 256$.

#### 5.2. The substitution model

In order to calculate the probability of emitting each of the 256 site patterns given a certain hidden state, we must choose a mutational model. In TRAILS, the substitution model is Jukes-Cantor [69], which assumes an equal mutation rate for all nucleotide changes, and thus predicts equal frequencies of 1/4 for the each of the four nucleotides at equilibrium. The resulting transition rate matrix $\boldsymbol{Q}$ is then parameterized by a single quantity $\mu$:

$$\boldsymbol{Q} = \frac{\mu}{4} \begin{pmatrix} -3 & 1 & 1 & 1 \\ 1 & -3 & 1 & 1 \\ 1 & 1 & -3 & 1 \\ 1 & 1 & 1 & -3 \end{pmatrix}.$$

The parameter $\mu$ is a re-scaling of the mutation rate $\nu$, such that $\mu = \frac{4}{3}\nu$. The transition probability matrix given at a certain time $u$ can therefore be computed as

$$\boldsymbol{P}(\mu, u) = e^{u\boldsymbol{Q}} = \frac{1}{4} + \frac{1}{4}e^{-u\mu} \begin{pmatrix} 3 & -1 & -1 & -1 \\ -1 & 3 & -1 & -1 \\ -1 & -1 & 3 & -1 \\ -1 & -1 & -1 & 3 \end{pmatrix}.$$

We can observe that there are only two possible transition probabilities, namely

$$P_{ij}(\mu, u) = \begin{cases} \frac{1}{4} + \frac{3}{4}e^{-u\mu}, & \text{if } i = j \\ \frac{1}{4} - \frac{1}{4}e^{-u\mu}, & \text{if } i \neq j \end{cases} \tag{6}$$

The above equation can thus be used to calculate the conditional probability of observing a certain nucleotide $b$ at time $u$, given a starting nucleotide $a$ at time 0, i.e., $\Pr(b \mid a)$.

#### 5.3. One coalescent event

As long as the genealogy of a hidden state follows a non-coalescing lineage, eq. (6) can be used to calculate the transition probability. However, there are parts of the hidden state genealogies that involve more than a single lineage, which happens at time intervals where coalescents between multiple lineages are allowed to happen. There are two different instances of this, either there is a single coalescent event, and two lineages merge into one (fig. S13), or there are two coalescent events in a single interval, and three lineages merge into one (fig. S14).

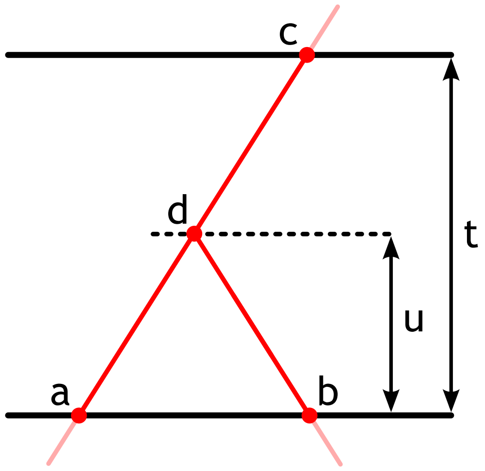

**Figure S13:** Diagram for an interval of length $t$ with a single coalescent event at time $u$. $a$, $b$, $c$ and $d$ represent nucleotides at different lineages and times.

For the case where a single coalescent happens, there are two initial lineages with nucleotides $a$ and $b$ at time 0, respectively, which they coalesce and segregate as a single lineage until observing a certain nucleotide $c$ at time $t$ (fig. S13). Let's assume the coalescent event happens at time $u$, such that we can record the observed nucleotide $d$ at the time of coalescent. By fixing the values of $a$, $b$, $c$, $d$ and $u$, we can calculate their joint probability given $a$ as

$$\Pr(b, c, d, u \mid a) = \left(e^{u\boldsymbol{Q}}\right)_{ad} \left(e^{u\boldsymbol{Q}}\right)_{bd} \left(e^{(t-u)\boldsymbol{Q}}\right)_{dc} \frac{e^{-u}}{\int_0^t e^{-v} dv}.$$

By integrating over all possible values of $u$, we get

$$\Pr(b, c, d \mid a) = \int_0^t \Pr(b, c, d, u \mid a) du = \frac{1}{1 - e^{-t}} \int_0^t e^{-u} \left(\frac{1}{4} + \alpha e^{-u\mu}\right) \left(\frac{1}{4} + \beta e^{-u\mu}\right) \left(\frac{1}{4} + \gamma e^{-(t-u)\mu}\right) du,$$

where

$$\alpha = \begin{cases} 3/4, & \text{if } a = d \\ -1/4, & \text{if } a \neq d \end{cases} \qquad \beta = \begin{cases} 3/4, & \text{if } b = d \\ -1/4, & \text{if } b \neq d \end{cases} \qquad \gamma = \begin{cases} 3/4, & \text{if } c = d \\ -1/4, & \text{if } c \neq d \end{cases}$$

This equation can be solved using Wolfram Mathematica [82] and can be consulted at `https://github.com/rivasiker/trails_paper/tree/main/emission_probability_formulas`.

Finally, we can calculate the joint probability of observing nucleotides $b$ and $c$ given $a$ by summing over all possibilities for $d$, i.e.,

$$\Pr(b, c \mid a) = \sum_d \Pr(b, c, d \mid a). \tag{7}$$

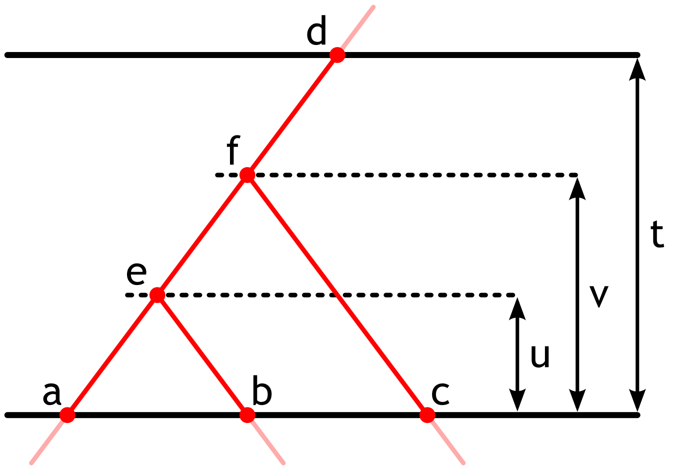

**Figure S14:** Diagram for an interval of length $t$ with a two coalescent events at times $u$ and $v$. $a$, $b$, $c$, $d$, $e$ and $f$ represent nucleotides at different lineages and times.

#### 5.4. Two coalescent events

Similarly, we can calculate the same formulas for when two coalescent events happen within the same interval. In this case, there are three different lineages at time 0, with nucleotides $a$, $b$ and $c$, respectively. The first coalescent event will happen at time $u$ between the $a$ and $b$ lineages, and afterwards, the second coalescent will happen at time $v$ with the remaining lineage $c$ (fig. S14). The single final lineage will last until time $t$. The nucleotide recorded at time $u$ is $e$, while that recorded at the time of the second coalescent $v$ is $f$.

Again, by fixing the values of $a$, $b$, $c$, $d$, $e$, $f$, $u$ and $v$, we can calculate their joint probability given $a$ as

$$\Pr(b,c,d,e,f,u,v \mid a) = \left(e^{u\boldsymbol{Q}}\right)_{ae} (e^{u}\boldsymbol{Q})_{eb} \left(e^{u\boldsymbol{Q}}\right)_{ef} \left(e^{(v+u)\boldsymbol{Q}}\right)_{fc} \left(e^{(t-v-u)\boldsymbol{Q}}\right)_{fd} \frac{3e^{-3u}e^{-(v-u)}}{k},$$

where $k = 1 + 0.5e^{-3t} - 1.5e^{-t}$, which is calculated from the convolution of two exponential distributions of rates 1 and 3. Similarly as before, we can integrate over all possible values of $v$ and $u$, such that

$$\Pr(b,c,d,e,f \mid a) = \int_0^t \int_u^t \Pr(b,c,d,e,f,u,v \mid a)\, dv\, du =$$
$$= \frac{3}{k} \int_0^t \int_0^v e^{-3u}e^{-(v-u)} \left(\frac{1}{4} + \alpha e^{-u\mu}\right) \left(\frac{1}{4} + \beta e^{-u\mu}\right) \left(\frac{1}{4} + \gamma e^{-(v-u)\mu}\right) \left(\frac{1}{4} + \delta e^{-v\mu}\right) \left(\frac{1}{4} + \epsilon e^{-(t-v)\mu}\right) du\, dv,$$

where

$$\alpha = \begin{cases} 3/4, & \text{if } a = e \\ -1/4, & \text{if } a \neq e \end{cases} \qquad \beta = \begin{cases} 3/4, & \text{if } b = e \\ -1/4, & \text{if } b \neq e \end{cases} \qquad \gamma = \begin{cases} 3/4, & \text{if } e = f \\ -1/4, & \text{if } e \neq f \end{cases}$$

$$\gamma = \begin{cases} 3/4, & \text{if } c = f \\ -1/4, & \text{if } c \neq f \end{cases} \qquad \epsilon = \begin{cases} 3/4, & \text{if } d = f \\ -1/4, & \text{if } d \neq f \end{cases}$$

This equation can also be solved using Wolfram Mathematica [82] and can also be consulted at `https://github.com/rivasiker/trails_paper/tree/main/emission_probability_formulas`.

Subsequently, we can calculate the join probability by summing over all values of $e$ and $f$:

$$\Pr(b,c,d \mid a) = \sum_e \sum_f \Pr(b,c,d,e,f \mid a). \tag{8}$$

#### 5.5. Piecing everything together

Using eqs. (6) to (8), we can calculate the Jukes-Cantor emission probabilities of all the hidden states. Each hidden state will have the three species involved in the ILS phenomenon (A, B and C), and an outgroup D, which is diverged enough from the rest of the species that no ILS can be observed. For all deep coalescent states ($V1_i^j$, $V2_i^j$ and $V3_i^j$) where the first and the second coalescent happen at the same time interval (i.e., $i = j$), then the joint probability for a certain combination of nucleotides (see labels in fig. S15 left) is given by

$$\Pr(a_0, a_1, b_0, b_1, c_0, c_1, abc_0, d_0) =$$

$$= \Pr(a_0) \cdot \Pr(a_1 \mid a_0) \cdot \Pr(b_1, c_1, abc_0, \mid a_1) \cdot \Pr(b_0 \mid b_1) \cdot \Pr(c_0 \mid c_1) \cdot \Pr(d_0 \mid abc_0),$$

Here, $\Pr(a_0)$ is the starting probability for observing each nucleotide, which in the Jukes-Cantor model corresponds to $1/4$, since all nucleotides are assumed to be equal in probability. Subsequently,

$$\Pr(a_0, b_0, c_0, d_0) = \sum_{a_1}\sum_{b_1}\sum_{c_1}\sum_{abc_0} \Pr(a_0, a_1, b_0, b_1, c_0, c_1, abc_0, d_0).$$

Alternatively, for all V0 states and for all deep coalescent states where the two coalescent events happen at different time intervals, then the joint probability for a combination of nucleotides (see labels in fig. S15 right) can be computed as

$$\Pr(a_0, a_1, b_0, b_1, ab_0, ab_1, c_0, c_1, abc_0, d_0) =$$

$$= \Pr(a_0) \cdot \Pr(a_1 \mid a_0) \cdot \Pr(b_1, ab_1 \mid a_1) \cdot \Pr(b_0 \mid b_1) \cdot \Pr(ab_0 \mid ab_1) \cdot \Pr(c_1, abc_0 \mid ab_1) \cdot \Pr(c_0 \mid c_1) \cdot \Pr(d_0 \mid abc_0),$$

and, subsequently,

$$\Pr(a_0, b_0, c_0, d_0) = \sum_{a_1}\sum_{b_1}\sum_{ab_0}\sum_{ab_1}\sum_{c_1}\sum_{abc_0} \Pr(a_0, a_1, b_0, b_1, ab_0, ab_1, c_0, c_1, abc_0, d_0).$$

All these formulas are similar in nature to Felsenstein's tree-pruning algorithm [83], but some of the conditional probabilities are calculated by integrating over all possible coalescent events rather than fixing the tree node to a certain value.

#### 6. Model parameterization and optimization

The transition and emission probabilities of TRAILS are parameterized by the speciation times, the ancestral effective population sizes, and the recombination rate. Given a sequence of observed states, these parameters are optimized so that the resulting model has the largest likelihood, thus performing parameter estimation.

Some of the parameters of the model are always fixed to ease the optimization. For example, the number of discretized time intervals between speciation events ($n_{\text{AB}}$) and the number of intervals in deep coalescence ($n_{\text{ABC}}$) are always kept fixed. On top of that, some other parameters are always optimized,

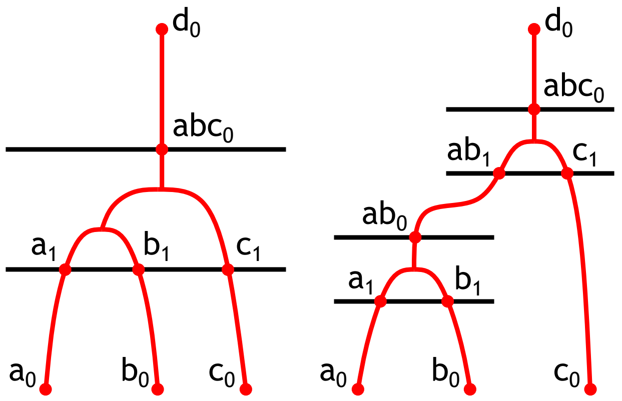

**Figure S15:** The two possible tree configurations for the emission probabilities, with labelled nucleotides at different time points.

such as the time between speciation events ($t_2$, in generations), the time from the end point of the second-to-last interval to the speciation time of the outgroup ($t_{\text{upper}}$, in generations), the (haploid) effective population size of the time between speciations and in deep coalescent ($N_{\text{AB}}$ and $N_{\text{ABC}}$, respectively), and the recombination rate ($r$, in crossovers per site per generation). We note that $t_{\text{upper}}$, which is only used to calculate the emission probabilities, provides an upper bound for coalescent events to occur between species A, B and C before they can coalesce with the outgroup. An assumption of the model is that the divergence between the outgroup and the rest of the species is large enough so that the majority of coalescent events have already happened before reaching the speciation event with the outgroup.

Additionally, we must specify the times from present to their corresponding speciation events. If the tree is ultrametric (in number of generations) and, thus, all lineages are sampled at time 0, then only one more parameter should be optimized, namely, the time from present to the first speciation event ($t_1$, in number of generations). This model is termed the *ultrametric* model. If instead each species has its own time, we would need one for each of the species, so the time from present to the first speciation event for A and B ($t_{\text{A}}$ and $t_{\text{B}}$, respectively), and the time from present to the second speciation event for C ($t_{\text{C}}$). The time from present to the third speciation event for the outgroup is calculated as $((t_{\text{A}} + t_{\text{B}})/2 + t_2 + t_{\text{C}})/2 + t_3$, where $t_3$ is the time from the second speciation event to the third speciation event, which can be calculated from $N_{\text{ABC}}$ and $t_{\text{upper}}$. This model is called the *non-ultrametric* model, and a diagram of it can be found in fig. 2A.

The mutation rate $\mu$ cannot be optimized together with the other parameters, because the same transition and emission probabilities can be computed by rescaling the parameters. In order to avoid this non-identifiability of the model, the mutation rate is fixed to a value of 1. Accordingly, all other parameters must be rescaled by the real mutation rate $\mu$, such that all times and effective population sizes are multiplied by the mutation rate ($t' = t\mu$ and $N' = N\mu$), and the recombination rate is divided by the mutation rate ($r' = r/\mu$).

The rescaled parameters are then optimized following a bound-constrained algorithm that can be user-specified (Nelder-Mead [74, 75], L-BFGS-B [45, 46], Powell [84], or TNC [85]), where the cost function is the log-likelihood of the model.

#### 7. Likelihood calculation and missing data

Following standard HMM theory [86, 87], we can use the forward algorithm to calculate the likelihood of the model given the data. To do so, we need to calculate first the transition probability matrix $\boldsymbol{A}$, the emission probability matrix $\boldsymbol{B}$, and the vector of starting probabilities $\boldsymbol{\pi}$. For a given sequence $\boldsymbol{o}$ of $n$ observations (in the case of TRAILS, the columns along a four-way genome alignment of length $n$), the forward algorithm can be implemented recursively as such:

1. Initialization: $\boldsymbol{\alpha}_1 = \boldsymbol{\pi}\boldsymbol{B}(o_1)$.
2. Recursion: $\boldsymbol{\alpha}_i = \boldsymbol{\alpha}_{i-1}\boldsymbol{A}\boldsymbol{B}(o_i)$, for $i \in (1, ..., n-1)$.
3. Termination: $L(\theta \mid \boldsymbol{o}) = P(\boldsymbol{o} \mid \theta) = \boldsymbol{\alpha}_n \boldsymbol{e}$.

Here, $\boldsymbol{\alpha}_i$ is the vector recording the probability of being at a certain hidden state after seeing the first $i$ observations, $\boldsymbol{B}(o_i)$ represents the column in the emission probability matrix corresponding to the state emitted at the $i$'th observation, and $\boldsymbol{e}$ is a column vector of ones of size $n$. In practice, this algorithm is implemented in the log-space to avoid underflow.

In the absence of missing data, all $o_i$ observations correspond to one of the 256 possible emitted states, and the forward algorithm can be applied as described above. If instead, $o_i$ contains missing information for all of species, i.e., if the observation is NNNN, the algorithm can be applied by substituting $\boldsymbol{B}(o_i)$ with $\boldsymbol{e}$.

Additionally, we can also observe partially missing data, where not all 4 sequences have missing information. For example, observations such as AAAN, ANAN, or ANNN are three of $5^4 - 4^4 - 1 = 368$ possible observations containing partially missing data. In such cases, each of the observations $o_i$ correspond to a set of emitted states $\boldsymbol{z}$, e.g., AAAN could only be AAAA, AAAC, AAAT or AAAG. Instead of substituting $\boldsymbol{B}(o_i)$ with $\boldsymbol{e}$ and losing information, we can instead substitute it with $\sum_j \boldsymbol{B}(z_j)$ for all $z_j \in o_i$. In the case of having only one missing nucleotide, there are only 4 possible $z_j$ states (as in the example above). For two missing nucleotides, there are $4^2 = 16$ possible $z_j$ states, and for three missing nucleotides, this number corresponds to $4^3 = 64$. In fact, the same formula can be used when all four nucleotides are missing, since $\sum_j \boldsymbol{B}(z_j)$ would correspond to summing over all of the columns ($4^4 = 256$) in the emission probability matrix $\boldsymbol{B}$, which yields a vector of ones.

In the current implementation of TRAILS, gaps are treated as missing data. It should be noted that a position in the alignment containing all gaps should be filtered out before using the sequence as input for the HMM.

#### 8. Implementation

All the functionality described here is implemented in a `python` package called `trails`, which can be downloaded and installed from pip (`https://pypi.org/project/trails-rivasiker`), and the source code can be browsed on GitHub (`https://github.com/rivasiker/trails`). The calculation of the transition and emission probability matrices is paralellized using `ray` [88]. Additionally, `pandas` [89] and `numpy` [73] are used for pre- and post-processing of the matrices. MAF alignment files are parsed using `biopython` [44]. Bound-constrained optimization algorithms are implemented in `SciPy` [90]. When possible, custom functions are njitted using `numba` [91] to boost performance.

Using the `trans_emiss_calc` from the `trails` package, one can calculate the transition and emission probability matrices in `python` by specifying the desired demographic parameters:

```python
from trails.optimizer import trans_emiss_calc
from trails.cutpoints import cutpoints_ABC

n_int_AB = 3
n_int_ABC = 3
mu = 2e-8

N_AB = 25000*2*mu
N_ABC = 25000*2*mu
t_1 = 240000*mu
t_2 = 40000*mu
t_3 = 800000*mu
t_upper = t_3-cutpoints_ABC(n_int_ABC,  1/N_ABC)[-2]
t_out = t_1+t_2+t_3+2*N_ABC
r = 1e-8/mu

transitions, emissions, starting, hidden_states, observed_states = trans_emiss_calc(
    t_1, t_1, t_1+t_2, t_2, t_upper, t_out,
    N_AB, N_ABC, r, n_int_AB, n_int_ABC)
```

If the alignment data is stored in a MAF file, then it can be parsed into `trails` using the `maf_parser` function:

```python
from trails.read_data import maf_parser
# The underlying species tree should be (((sp1,sp2),sp3),out);
# These names should match the species names in the MAF file
sp_lst = ['sp1', 'sp2', 'sp3', 'out']
observed = maf_parser('chr1.maf', sp_lst)
```

If the alignment data is split up into different alignment blocks in the MAF file, the resulting parsed alignment will also contain all those split blocks.

We can calculate the likelihood of the model given the data using the `loglik_wrapper` function:

```python
from trails.optimizer import loglik_wrapper
loglik = loglik_wrapper(transitions, emissions, starting, observed)
```

If the data is split into several alignment blocks, then the log-likelihood will be calculated independently for each block, i.e., the HMM will be re-initialized in each block. The overall log-likelihood (the cost function) will be the sum of the log-likelihood of all the blocks.

Finally, the `optimizer` function conveniently wraps all this together to perform numerical optimization given the data and using a specified bound-constrained algorithm. The bounds are specified using a `python` dictionary, which contains a list for each of the optimized parameters with the initial value, the lower bound and the upper bound. This dictionary should be supplied in the `optim_params` argument. Moreover, the fixed parameters, which in this case are the number of intervals between the two speciation events ($n_{\mathrm{AB}}$) and the number of intervals deep in time ($n_{\mathrm{ABC}}$), are also specified using a dictionary, and supplied in the `fixed_params` argument. The results are saved on a csv file specified by the path in `res_name`. Finally, the optimization method is specified using the `method` argument. All together:

```python
dct = {
    't_1':     [t_1,     t_1/10,     t_1*10],
    't_2':     [t_2,     t_2/10,     t_2*10],
    't_upper': [t_upper, t_upper/10, t_upper*10],
    'N_AB':    [N_AB,    N_AB/10,    N_AB*10],
    'N_ABC':   [N_ABC,   N_ABC/10,   N_ABC*10],
    'r':       [r,       r/10,       r*10]
    }
dct2 = {'n_int_AB':n_int_AB, 'n_int_ABC':n_int_ABC}
res = optimizer(
    optim_params = dct,
    fixed_params = dct2,
    V_lst = observed,
    res_name = 'results.csv',
    method = 'L-BFGS-B'
    )
```

Alternatively, if the speciation tree is not ultrametric, the dictionary for `optim_param` can also contain independent speciation times for species A, B, and C, in which case all those will be optimized:

```
dct = {
    't_A':      [t_1,      t_1/10,       t_1*10],
    't_B':      [t_1,      t_1/10,       t_1*10],
    't_C':      [t_1+t_2,  (t_1+t_2)/10, (t_1+t_2)*10],
    't_2':      [t_2,      t_2/10,       t_2*10],
    't_upper': [t_upper, t_upper/10,   t_upper*10],
    'N_AB':     [N_AB,     N_AB/10,      N_AB*10],
    'N_ABC':    [N_ABC,    N_ABC/10,     N_ABC*10],
    'r':        [r,        r/10,         r*10]
    }
dct2 = {'n_int_AB':n_int_AB, 'n_int_ABC':n_int_ABC}
res = optimizer(
    optim_params = dct,
    fixed_params = dct2,
    V_lst = observed,
    res_name = 'results.csv',
    method = 'L-BFGS-B'
    )
```

The underlying functions are parallelized, so multiple cores will make the calculations faster, especially when $n_{\mathrm{ABC}}$ gets larger.

#### 9. Runtime

As the number of discretized time intervals between speciation events ($n_{\mathrm{AB}}$) increases and, more importantly, as the number of discretized time intervals in deep coalescence ($n_{\mathrm{ABC}}$) increases, the model becomes more complex. This complexity is reflected in the number of hidden states, which amounts to $n_{\mathrm{AB}} \times n_{\mathrm{ABC}} + 3 \times \left[n_{\mathrm{ABC}} + \binom{n_{\mathrm{ABC}}}{2}\right]$, and it heavily impacts the runtime of each optimization round. The optimization procedure requires two computationally expensive steps. The first step involves calculating the transition probability matrix given a certain demographic model, $n_{\mathrm{AB}}$ and $n_{\mathrm{ABC}}$. Afterward, the second step corresponds to calculating the likelihood of the model given some data. While this second step is the most computationally heavy for small values of $n_{\mathrm{ABC}}$, it quickly becomes surpassed by the time spent calculating the transition probability matrix (fig. S16). Conversely, posterior decoding of a fitted model takes around the same time as computing the likelihood.

Based on the simulations used for fig. 2, around 150-to-200 iterations of the optimization procedure were necessary to achieve convergence. Given the runtimes of fig. S16, for each iteration when $n_{\mathrm{AB}} = n_{\mathrm{ABC}} = 5$, TRAILS spends $\sim 170$ seconds for computing the transition probability matrix, and $\sim 75$

seconds for the likelihood calculation for a 10Mb region. For a total of 150 iterations, this amounts to $\sim$ 10 hours of computations to achieve convergence.

#### 10. Parametric bootstrapping

To quantify the variability of the parameters estimated by TRAILS, 20 genomes, 50 Mb in length, were simulated from the model fitted with the parameters estimated for the HCGO alignment. After performing parameter optimization using TRAILS, the 20 estimates were used to fit a normal distribution per parameter (fig. S18), and 95% confidence intervals were computed from the fitted normal distribution (table S2).

| Parameter | Estimated value | Bootstrapped mean | Bootstrapped SD | Low CI | High CI |
|---:|---:|---:|---:|---:|---:|
| H-to-HC | 5.51 | 5.49 | $3.08 \times 10^{-2}$ | 5.43 | 5.54 |
| C-to-HC | 5.84 | 5.83 | $2.87 \times 10^{-2}$ | 5.77 | 5.87 |
| G-to-HCG | 11.70 | 11.71 | $2.31 \times 10^{-2}$ | 11.66 | 11.74 |
| O-to-HCGO | 19.28 | 19.31 | $4.60 \times 10^{-2}$ | 19.22 | 19.38 |
| HC-to-HCG | 4.89 | 4.90 | $3.05 \times 10^{-2}$ | 4.84 | 4.95 |
| HCG-to-HCGO | 8.15 | 8.18 | $3.99 \times 10^{-2}$ | 8.10 | 8.24 |
| $N_{\mathrm{AB}}$ | 167400 | 168323 | 1416 | 165548 | 170361 |
| $N_{\mathrm{ABC}}$ | 101290 | 101058 | 302 | 100467 | 101492 |
| $\rho$ | $1.191 \times 10^{-8}$ | $1.189 \times 10^{-8}$ | $8.361 \times 10^{-11}$ | $1.172 \times 10^{-8}$ | $1.200 \times 10^{-8}$ |

**Table S2:** Estimated values, and mean, standard deviation (SD) and 95% confidence intervals (CI) of normal distributions fitted on the parametric bootstrap replicates.

The same 20 replicates were also used to explore how parameters covary when estimated. Results show that several pairs of parameters are correlated (fig. S19). For example, both $t_{\mathrm{A}}$ and $t_{\mathrm{B}}$ are significantly negatively correlated with $t_2$, since in order to compensate for a large value of $t_2$, the lengths of the tip branches need to decrease. On the other hand, $t_2$ is positively correlated with $N_{\mathrm{AB}}$, since, in order to keep the same level of ILS, $t_2$ and $N_{\mathrm{AB}}$ must increase or decrease by the same factor, since $\mathrm{ILS} = \frac{2}{3}\exp(-t_2/(2N_{\mathrm{AB}}))$. Also, notably, $\rho$ is not significantly correlated to any of the parameters.

#### 11. Bibliography

List of references for the Supporting Information is the same as for the main text.

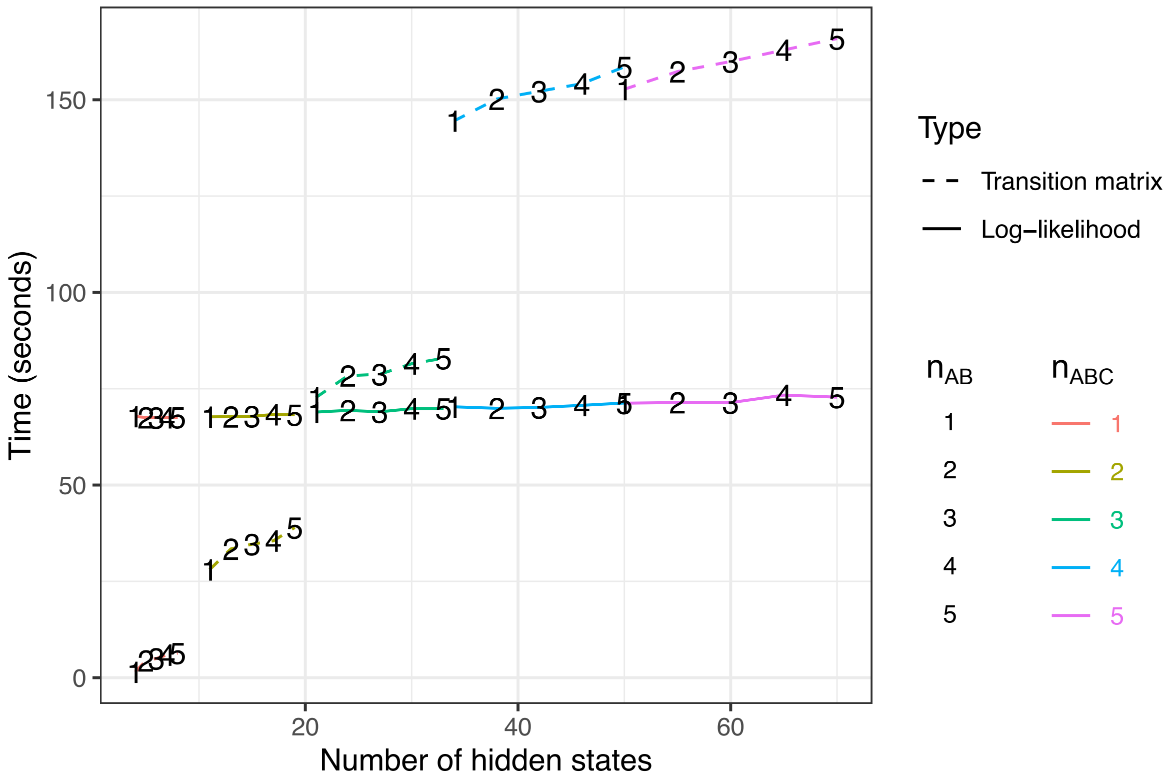

**Figure S16:** Runtime for calculating the transition probability matrix and the log-likelihood for a 1-Mb region of the HCGO alignment using the current implementation of TRAILS. The data points are averaged over 5 runs per combination of $n_{AB} \in [1, 2, 3, 4, 5]$ and $n_{ABC} \in [1, 2, 3, 4, 5]$. All these computations were performed on an 8-core MacBook Pro with Intel Core i5 processor and 8 GB or memory.

.

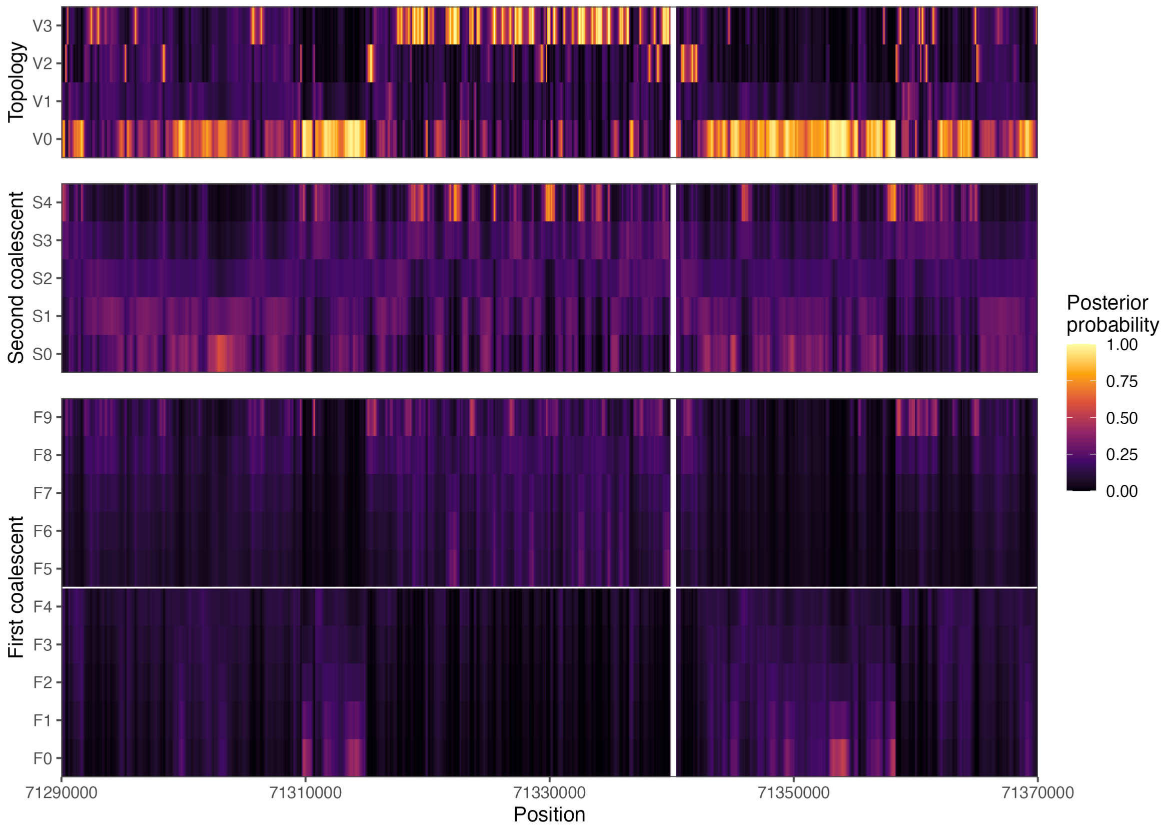

**Figure S17:** Posterior decoding of the topology and coalescent times of a region in chromosome 1 showing an excess of V3 topology.

.

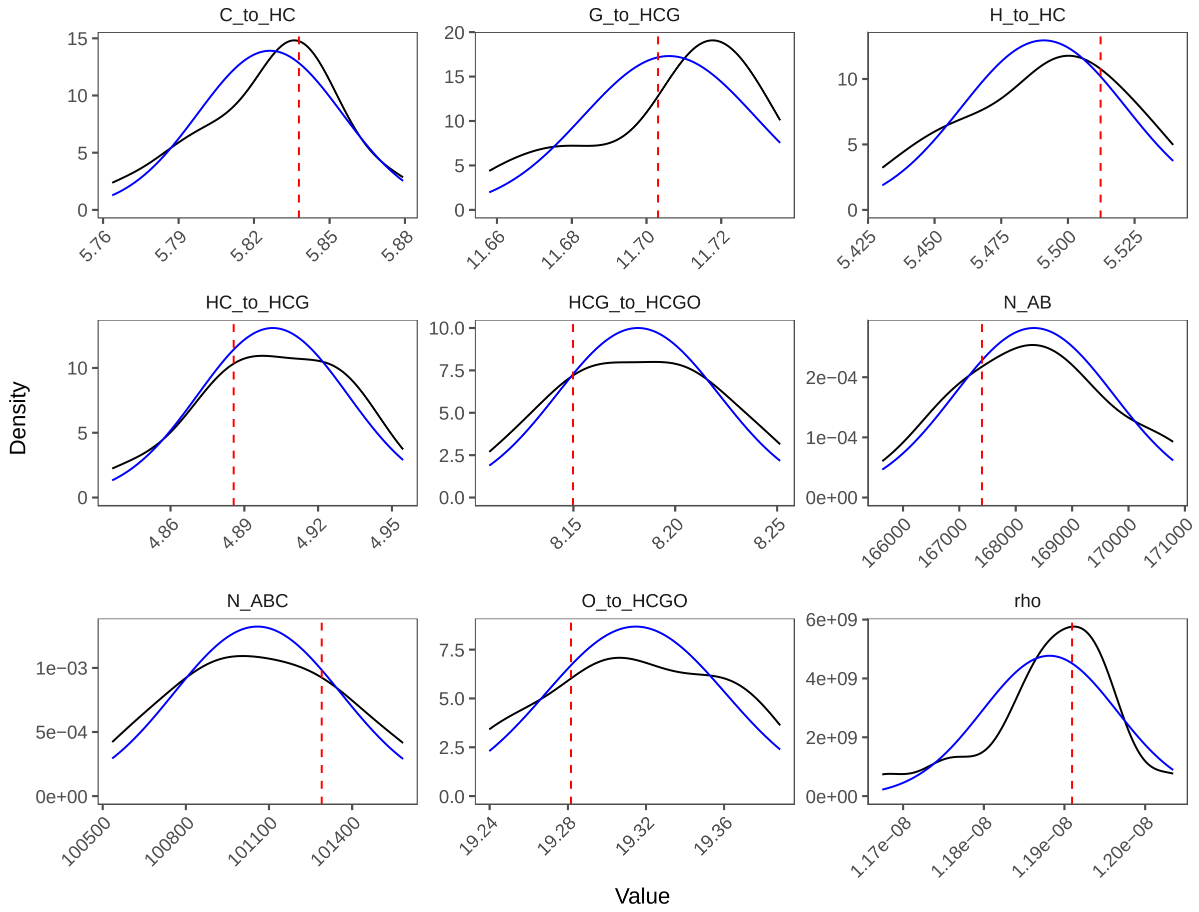

**Figure S18:** Kernel densities (black) and fitted normal distributions (blue) for the parametric bootstrap replicates. True values are plotted as vertical dashed red lines.

.

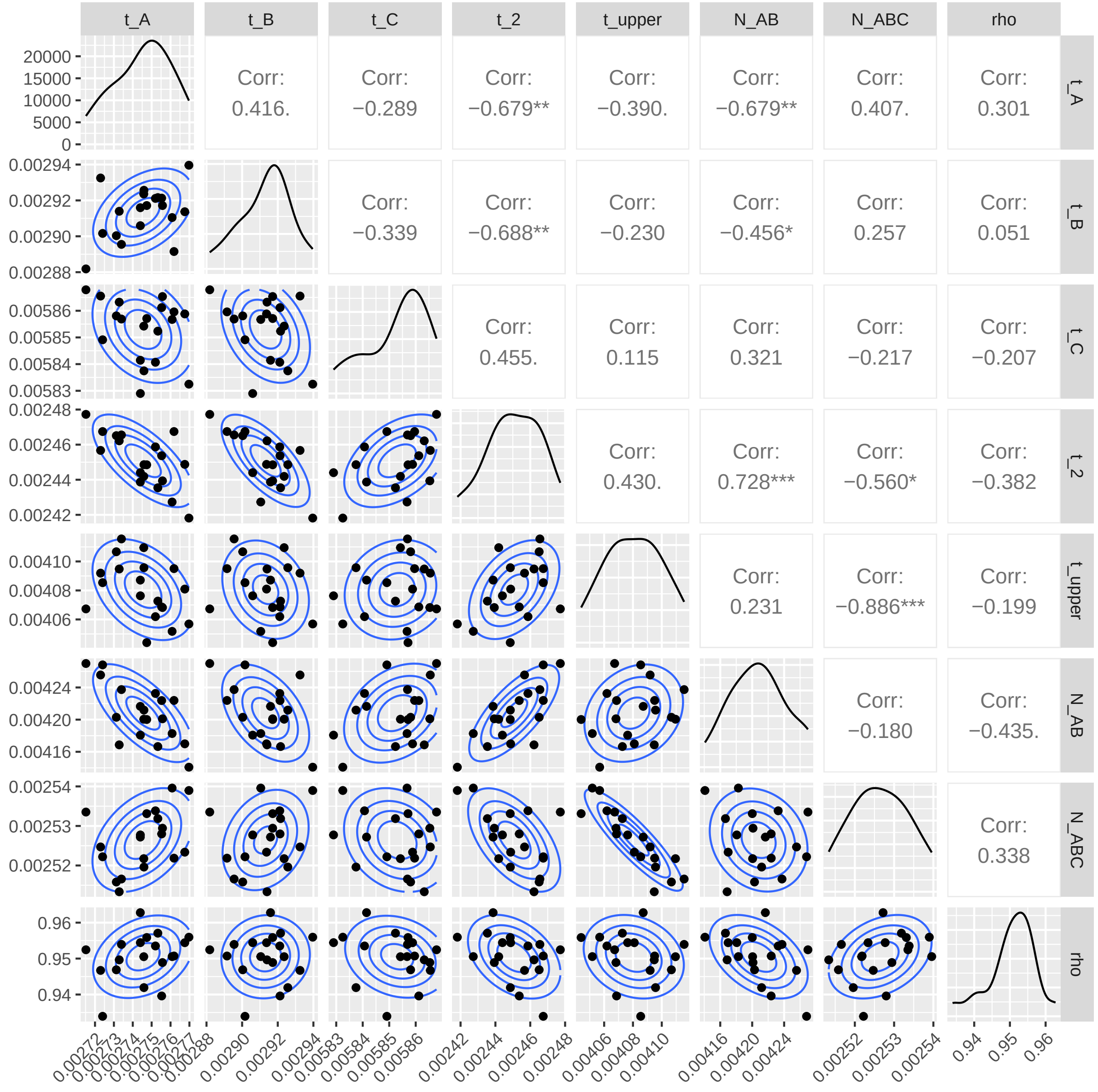

**Figure S19:** Pairwise correlation analysis for the raw values of the estimates of the parametric bootstrapping. The kernel densities are plotted in the diagonal. The upper diagonal contains correlation coefficients, with significance levels of <0.001 ("***"), <0.01 ("**"), <0.05 ("*"), and <0.1 ("."). The lower diagonal contains pairwise scatter plots, with contour lines corresponding to the density of a fitted bivariate normal distribution.
